# DNA-guided transcription factor cooperativity shapes face and limb mesenchyme

**DOI:** 10.1101/2023.05.29.541540

**Authors:** Seungsoo Kim, Ekaterina Morgunova, Sahin Naqvi, Maram Bader, Mervenaz Koska, Alexander Popov, Christy Luong, Angela Pogson, Peter Claes, Jussi Taipale, Joanna Wysocka

**Author notes:** These authors contributed equally (Ekaterina Morgunova, Sahin Naqvi).

## Abstract

Transcription factors (TFs) can define distinct cellular identities despite nearly identical DNA-binding specificities. One mechanism for achieving regulatory specificity is DNA-guided TF cooperativity. Although *in vitro* studies suggest it may be common, examples of such cooperativity remain scarce in cellular contexts. Here, we demonstrate how ‘Coordinator’, a long DNA motif comprised of common motifs bound by many basic helix-loop-helix (bHLH) and homeodomain (HD) TFs, uniquely defines regulatory regions of embryonic face and limb mesenchyme. Coordinator guides cooperative and selective binding between the bHLH family mesenchymal regulator TWIST1 and a collective of HD factors associated with regional identities in the face and limb. TWIST1 is required for HD binding and open chromatin at Coordinator sites, while HD factors stabilize TWIST1 occupancy at Coordinator and titrate it away from HD-independent sites. This cooperativity results in shared regulation of genes involved in cell-type and positional identities, and ultimately shapes facial morphology and evolution.

## Introduction

Sequence-specific transcription factors (TFs) play a key role in controlling gene expression programs during embryonic development and in response to environmental cues. Forcibly expressing certain TFs often referred to as master regulators is sufficient to change a cell’s identity^1^. For example, overexpression of MyoD1 can convert fibroblasts into myoblasts^2,3^, while overexpression of NeuroD1 can induce terminal neuron differentiation^4^. TFs bind specific, typically short DNA sequences called motifs, and recruit coactivators or corepressors to modulate transcription^5,6^. However, many TFs fall into large families, and share highly conserved DNA-binding domains (DBDs) that often bind very similar DNA motifs across family members^3,6^. Even TFs that drive divergent gene expression programs can bind the same motifs^3^, raising the question of how they achieve regulatory specificity. Among the largest TF families in humans are homeodomain (HD) (with >200 TFs) and basic helix-loop-helix (bHLH, >100 TFs) proteins, which are well known for prominent roles in driving positional identities (e.g. the HOX genes that specify anterior-posterior segmental identity^7^) and cell type identities^8^ (e.g. MyoD1 and NeuroD1), respectively. And yet, within each TF family there is extensive similarity of the DNA binding motifs among members. Most bHLH factors recognize a subset of CANNTG sequences collectively called the ‘E-box’^9^ (each bHLH subfamily prefers a certain variant such as CAgaTG but can often also bind other variants), while the motif TAATT[A/G] is bound by roughly a third of all HD TFs in humans^10,11^.

Cooperative TF binding is a potential mechanism for achieving DNA-binding specificity among the various TFs of large DBD families and for integration of multiple biological inputs at *cis*-regulatory elements. Diverse mechanisms underlying TF cooperativity have been described in the literature^12^. In the simplest and most well-understood mechanism, direct protein-protein contacts mediate stable TF hetero- or homo- dimer formation in solution, which in turn allows for stronger DNA binding (and regulatory activity) by the resulting TF complex. Another, less specific, mechanism is nucleosome-mediated cooperativity, in which essentially any TFs with binding sites within ∼150 bp can indirectly cooperate by competing with the same nucleosome; this type of cooperativity does not require stereotypical spacing or organization of the motifs, sometimes referred to as ‘motif grammar’^13^. Finally, certain TFs can cooperatively bind juxtaposed DNA sites arranged in specific orientation and distance without forming stable, direct protein-protein interactions; we will refer to this form of co-binding as ‘DNA-guided cooperativity’. *In vitro* analysis of TF pairs using consecutive affinity-purification systematic evolution of ligands by exponential enrichment (CAP-SELEX) revealed that DNA-guided cooperativity may be a common feature of gene regulation^14^. However, beyond *in vitro* work, most cellular studies of this mechanism and its biological function have been limited to few well-documented examples, such as the pluripotency factors OCT4 and SOX2, which bind at a composite motif that combines the two TFs’ individual motifs^14^ to facilitate chromatin opening^15,16^. In other cases, composite motifs have been observed in DNA sequence analyses of enhancers^17–19^, but the mechanisms of cooperativity and selectivity among related TFs binding at these sites remain unexplored.

We previously serendipitously discovered a novel, 17-bp long DNA sequence motif with strong evidence of endogenous cellular function, which we termed ‘Coordinator’^20^. By comparing enhancer landscapes in human and chimpanzee facial progenitor cells called cranial neural crest cells (CNCCs) and analyzing the underlying DNA sequence changes, we uncovered motifs whose gains and losses correlated with changes in enhancer activity, as measured by chromatin marks such as histone 3 lysine 27 acetylation (H3K27ac) and DNA accessibility (**Figure 1A**). The Coordinator motif, discovered through *de novo* sequence analysis, was far more predictive of active chromatin states and species bias in enhancer activity than any known TF motif^20^. DNA sequence substitutions between the human and chimp genomes that resulted in gains of fit to the Coordinator consensus motif in either species were associated with gains in enhancer activity in that same species. We therefore hypothesized that the *trans*-regulatory complex that recognizes the motif plays an outsized role in coordinating enhancer activity in CNCCs, and hence named the motif Coordinator. Although the motif was not previously annotated to a known regulatory complex, it did not escape our attention that parts of Coordinator contain the TAATT[A/G] motif bound by many HD factors and a version of the CANNTG ‘E-box’ motif bound by most bHLH factors, separated by a fixed spacing (**Figure 1A**). Given the large number of bHLH and HD factors encoded by the human genome^6^, the Coordinator motif represents a valuable opportunity to gain insight into the mechanisms of specificity and functional implications of TF co-binding in a biologically relevant context.

**Figure 1.**
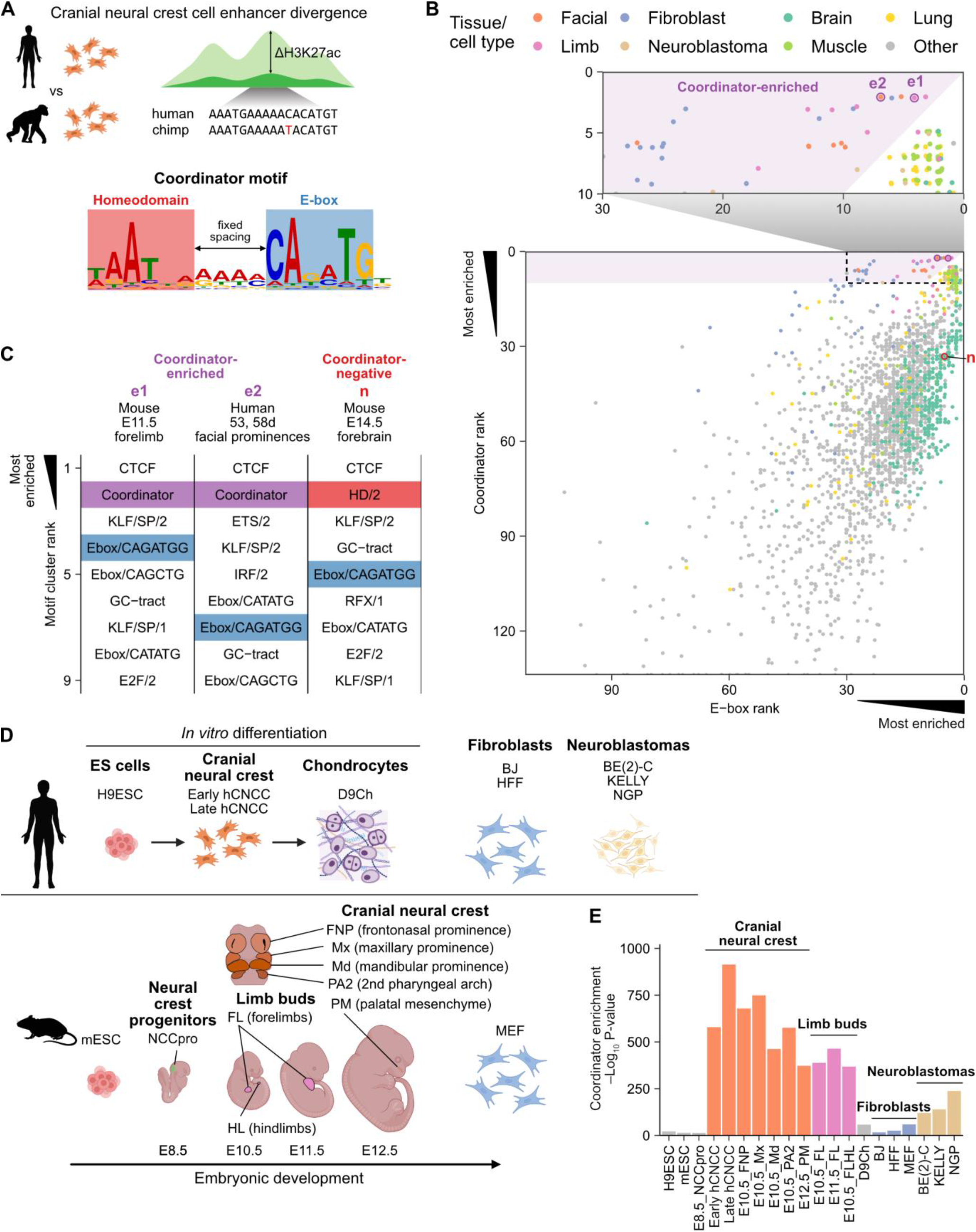
The ‘Coordinator’ motif is active specifically in embryonic face and limb mesenchyme. **A**. Schematic of the ‘Coordinator’ motif and its discovery from studying human and chimpanzee cranial neural crest cell enhancer divergence. **B**. Rankings of Coordinator and its constituent Ebox/CAGATGG motif in enrichment in the top 10,000 distal accessible regions, for all DNase-seq and ATAC-seq datasets on ENCODE. e1, e2, and n indicate examples of Coordinator-enriched and Coordinator-negative samples, detailed in (**C**). Points are jittered to avoid overplotting. Zoom-in highlights the prevalence of facial, limb, and fibroblast samples among those with Coordinator motif enrichment (Coordinator rank < 10 and Coordinator rank < E-box rank). **C**. Examples of Coordinator-enriched and Coordinator-negative samples, with their top 9 motif clusters. Coordinator is highlighted in purple, the best E-box motif match to Coordinator (Ebox/CAGATGG) in blue, and the best homeodomain motif match to Coordinator (HD/2) in red. **D**. Schematic of cell types and tissues analyzed in (**E**). **E**. Coordinator motif enrichment p-value by AME^99^ (analysis of motif enrichment), with cranial neural crest samples in orange, limb buds in pink, fibroblasts in blue, and neuroblastomas in tan (others in gray).

Here, we sought to systematically identify TFs that bind the Coordinator motif, determine their molecular functions in an endogenous cellular context, and dissect the mechanisms underlying their cooperativity and selectivity. We show that despite the abundance of cell types with high enrichment of the HD and E-box motifs in their active regulatory regions, the Coordinator motif is specific to the developing face and limb mesenchyme. We find that Coordinator is co-bound by a bHLH factor TWIST1, a driver of mesenchymal identity, and by HD TFs associated with defined regional identities in the face and limb, including ALX1, ALX4, MSX1, and PRRX1, all linked to craniofacial, and in some cases also limb, congenital anomalies in humans^21–24^. Using endogenous degron- and epitope-tagging and nonsense mutations, we studied these TFs’ genomic binding and function in human CNCCs (hCNCCs). We found that TWIST1 is required for accessibility, HD TF binding, and enhancer activity at Coordinator sites. In turn, the HD TFs primarily (and at least in part redundantly) function to stabilize TWIST1 binding at Coordinator sites and to titrate it away from its canonical and HD-independent E-box and double E-box sites towards Coordinator. This co-binding at Coordinator sites drives shared transcriptional functions, with genes associated with lineage and regional identities in the facial mesenchyme showing dependency on both TWIST1 and the ALX factors. TWIST1 and HD TFs do not interact in solution, but instead their cooperativity is guided by the Coordinator motif DNA sequence. We solve a crystal structure of TWIST1, its heterodimerization partner TCF4, and ALX4, bound to the Coordinator DNA, revealing the structural basis of this DNA-guided cooperativity. Finally, we show that genetic variants at loci encoding Coordinator-binding TFs are associated with overlapping aspects of the normal-range facial variation in humans, and that *cis*-regulatory regions dependent on these TFs are enriched for facial shape heritability. Together, our work illustrates how DNA-guided TF cooperativity can facilitate regulatory specificity and integrate cell lineage and positional information to shape the development and evolution of the face and limb mesenchyme.

## Results

### The ‘Coordinator’ motif is active specifically in embryonic face and limb mesenchyme

The Coordinator motif^20^ is a composite of the TAATT[A/G] motif bound by many HD factors and a version of the CANNTG E-box bound by most bHLH factors^6^, separated by a fixed spacing (**Figure 1A**). Therefore, we wondered whether any cell types other than CNCCs also exhibit enrichment for the Coordinator motif in their active *cis*-regulatory regions. We first defined a robust signature of Coordinator activity, based on the observation that in hCNCCs the Coordinator motif is enriched most highly in the top ∼10,000 promoter-distal open chromatin peaks as defined by the assay for transposase-accessible chromatin with sequencing (ATAC-seq) (**Figure S1A**). We searched for enrichment of known TF motifs in the top 10,000 distal open chromatin regions for each of the 549 ATAC-seq and 1,781 DNase-seq datasets from human and mouse in ENCODE^25^, and then collapsed similar motifs into 287 motif clusters to remove redundant motifs (**Materials and Methods**). Finally, we checked the ranking of the Coordinator motif cluster and the motif clusters best matching its constituent motifs, the Ebox/CAGATGG and HD/2 clusters, and considered a sample to exhibit Coordinator activity when: 1) Coordinator is among the top 10 motif clusters, and 2) Coordinator is ranked higher than either constituent motif cluster (**Figure 1B,C**). This approach can also be applied to other known composite motifs such as the OCT:SOX motif, which meets similar criteria in pluripotent stem cells (**Figure S1B**). By using only the strongest peaks and motif rankings, this approach should be more robust to variation in sample quality and cellular heterogeneity than approaches using all open chromatin regions or quantitative metrics of motif enrichment.

As expected, the embryonic facial prominence samples, which are largely comprised of CNCCs, exhibit strong Coordinator motif activity, with Coordinator ranked as the second most-enriched motif, after CTCF (**Figure 1B,C**). However, many developing limb samples and a smaller subset of fibroblast and neuroblastoma samples also meet our definition of Coordinator activity. Notably, neuroblastoma is a cancer originating from neural crest-derived lineages^26^, whereas fibroblasts are mesenchymal cells of either mesodermal or neural crest origin^27^. Importantly, across analyzed cell types and tissues, most samples showing strong enrichment of E-box and HD motifs are not enriched for Coordinator (**Figure 1B, Figure S1C**). For example, many developing brain samples are highly enriched in both E-box and HD halves of the motif but not for the combined motif (**Figure 1B,C; Figure S1C**). To corroborate this finding, we gathered additional published human and mouse ATAC-seq datasets from cell types related to those in which we initially detected Coordinator enrichment and from relevant negative controls (**Figure 1D**)^28–39^. We plotted the p-values of Coordinator motif enrichment and found that *in vitro*-derived mesenchymal hCNCCs and mouse embryonic facial prominences of CNCC origin have the strongest enrichment, followed closely by limb bud samples, with much lower enrichment in neuroblastomas and fibroblasts (**Figure 1E**). Altogether, these results indicate that the Coordinator motif is selectively enriched in the accessible *cis*-regulatory regions of the developing face and limb mesenchyme.

### TWIST1 binds Coordinator across tissues with diverse homeodomain TF expression

Due to the discovery of Coordinator from *de novo* motif analysis and the similarity of binding motifs for TFs among the bHLH and HD families, the identity and number of TFs that bind Coordinator remains unclear. To systematically assess candidate Coordinator-binding TFs, we searched for TFs that have: 1) binding motifs consistent with the constituent E-box or HD halves of Coordinator, and 2) high expression levels specifically in cell types with Coordinator enrichment in open chromatin.

First, we aligned known TF motifs [derived from either chromatin immunoprecipitation DNA sequencing (ChIP-seq) data or *in vitro* specificity measurements using SELEX or protein binding microarrays (PBM), as indicated] against each half of the Coordinator motif, beginning with the E-box typically bound by bHLH factors (**Figure 2A; Figure S2A**). A total of 54 TFs have motifs that aligned to the E-box, of them 40 in the bHLH family. TWIST1 is the only TF with a known motif that spans both the E-box and the HD motif, though this motif is derived from ChIP-seq in neuroblastoma cells^39^ and as a bHLH factor, it only directly binds the E-box. In fact, previously published ChIP-seq for TWIST1 overexpressed in human mammary epithelial cells revealed binding to the single or double E-box motifs^40^. Next, we examined the RNA levels of each candidate TF and their correlation with Coordinator motif enrichment p-values in active regulatory regions across cell types. TWIST1 was the TF with the highest correlation (**Figure S2B**). Remarkably, its expression levels were nearly perfectly correlated with Coordinator motif enrichment (r = 0.934; **Figure S2C**). Indeed, our previous work detected Coordinator motif enrichment at TWIST1 ChIP-seq peaks from hCNCCs^28^. To confirm that TWIST1 binds Coordinator *in vivo* in cell types with enrichment for Coordinator in active regulatory regions, we performed ChIP-seq for Twist1 in dissected E10.5 mouse embryos (**Figure 2B**), separately testing the frontonasal prominences (FNP), maxillary prominences (Mx), mandibular prominences (Md), forelimbs (FL), and hindlimbs (HL). We compared these to ChIPs from hCNCCs and previously published data from the neuroblastoma cell lines BE(2)-C and SHEP21^39^. Across all of these cellular contexts, most of the strongest TWIST1 peaks contained the Coordinator motif, but weaker peaks were progressively less likely to contain the Coordinator motif (using a threshold of p < 10^-4^). However, compared to the hCNCCs, facial prominences, and limb buds, which sustained high Coordinator motif frequencies (>50%) for the top 20,000 peaks or more, neuroblastomas only had such motif frequencies in the top few thousand peaks (despite a greater total number of peaks). This rapid falloff is consistent with the weaker Coordinator enrichment in neuroblastoma open chromatin (**Figure 1E**).

**Figure 2.**
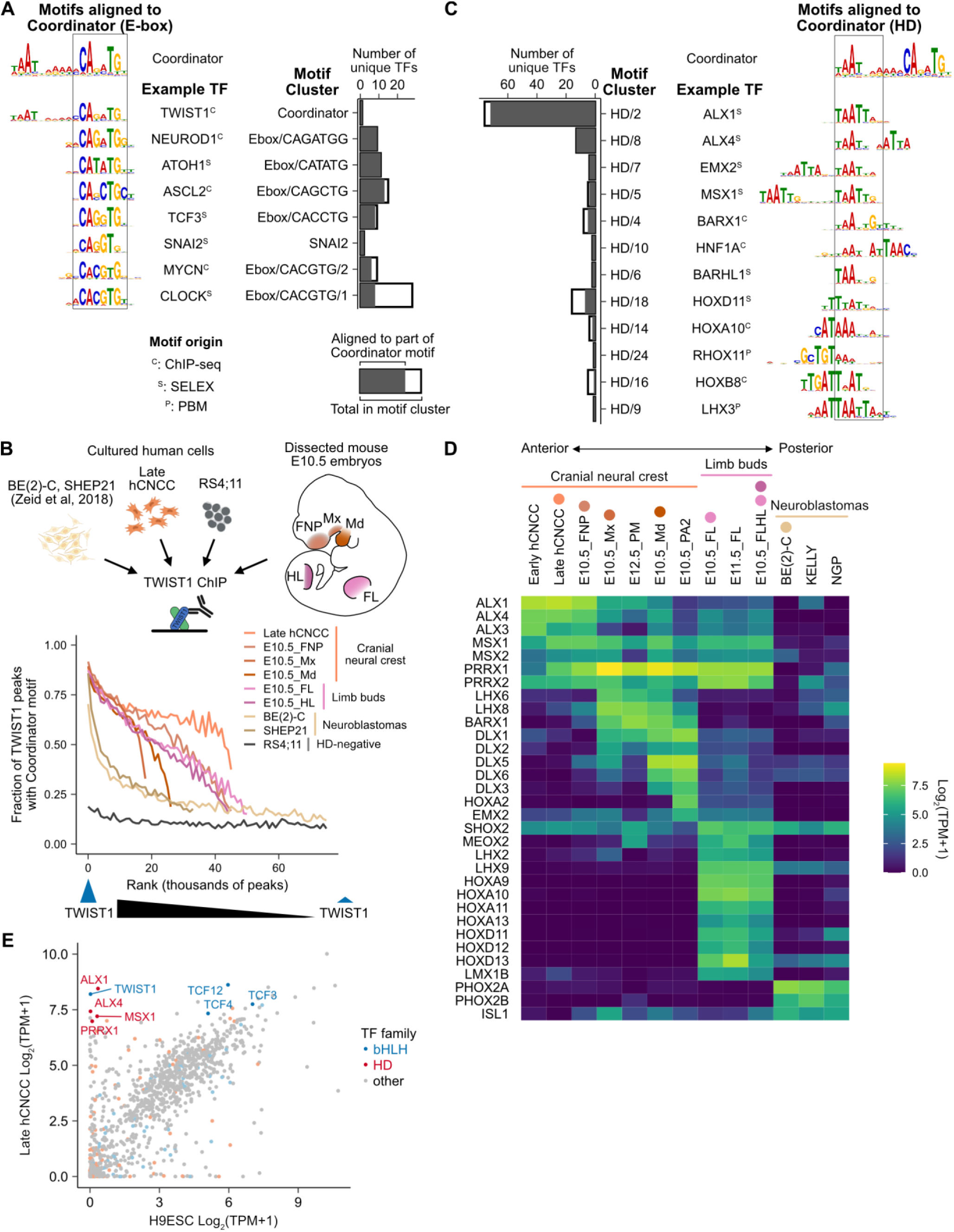
TWIST1 binds Coordinator across tissues with diverse homeodomain TF expression. **A**. Motif clusters with motifs aligned to the E-box portion of Coordinator (marked by box) with logos and TF names of examples, and the total number of motifs in each cluster (black outline) and the number that successfully aligned to the Coordinator portion (filled in gray, TOMTOM q-value < 0.4). TF name superscript indicates origin of motif: C, ChIP; P, PBM; S, SELEX. **B**. TWIST1 ChIP-seq analysis in the indicated human cell types and mouse tissues. Schematic of E10.5 mouse embryo illustrates the dissected regions. The strongest TWIST1 peaks and those in cell types with high HD TF activity tend to contain a Coordinator motif (p < 1e-4, within 100 bp of the summit). TWIST1 ChIP-seq peaks are ranked from strongest to weakest (left to right) and grouped into bins of 1000 peaks. **C**. As in (**A**), but for the homeodomain (HD) portion of the Coordinator motif. **D**. HD TF expression varies across cell/tissue types with Coordinator enrichment. RNA expression is shown for all HDs with motifs aligning to Coordinator and appreciable expression in at least one cell type. Colored circles above cell type names correspond to the schematic and lines in (**C**). **E**. TWIST1 and multiple HDs are specifically highly expressed in human cranial neural crest cells (hCNCC) but not H9 embryonic stem cells (H9ESC). bHLH and HD family TFs are highlighted in blue and red, respectively (others in gray). TFs with high expression specifically in hCNCC are in darker colors and labeled with their names.

Next, we focused on candidate factors binding the HD portion of Coordinator. Of the 129 TFs with motifs aligned to the HD half of Coordinator, 108 contain an HD DNA-binding domain (**Figure 2C**; **Figure S2D**). Of these, 32 are expressed moderately or highly in at least one cell type with Coordinator enrichment (including cranial neural crest, limb buds, and neuroblastomas) (**Figure 2D**). However, no single clear candidate emerged that was expressed in all Coordinator-positive cell types and could explain the quantitative variation in Coordinator activity. Instead, every cell type expresses multiple HD TFs robustly, with groups of HDs showing overlapping expression in distinct regions of the developing face and limbs, consistent with their previously described association with specific positional identities^41–43^ (such as along the anterior-posterior axis). For example and as expected, the ALX factors are enriched in the frontonasal prominence, which gives rise to the upper part of the face, whereas DLX factors are enriched in the mandibular prominence, which gives rise to the lower jaw, while the posterior HOX genes are expressed in the limb buds but not the face (**Figure 2D**). In contrast, neuroblastoma cells express known neuronal/glial regulators PHOX2A/B and ISL1 highly but lack most mesenchymally expressed HD factors.

To test whether these HDs collectively enable TWIST1 binding to Coordinator, we searched the Cancer Cell Line Encyclopedia^44^ (CCLE) for any cell lines that have high RNA levels of TWIST1 but minimal levels of these candidate Coordinator-binding HDs (**Figure S2E**). One of the best matches to our criteria was RS4;11, an acute lymphoblastic leukemia cell line with a t(4;11) translocation. We then performed TWIST1 ChIP-seq in RS4;11 cells and found that although TWIST1 binds DNA robustly in RS4;11 cells, those sites are enriched for the single and double E-box motifs (**Figure S2F**), similar to the motifs bound by TWIST1 upon overexpression in human mammary epithelial cells^40^. In contrast, the Coordinator motif frequency (< 20%) is around that of the weakest TWIST1 peaks in any context (**Figure 2B**). These results suggest that TWIST1 binds Coordinator only in cell types in which certain HD proteins are also present.

### Multiple homeodomains co-bind Coordinator motif with TWIST1

To study the mechanisms and functional role of TWIST1 cooperation with HD TFs at Coordinator in a biologically relevant context, we turned to our previously characterized *in vitro* model of human embryonic stem cell (hESC) differentiation to hCNCCs^20,28,45,46^. TWIST1 is the only bHLH TF selectively expressed in hCNCCs compared to hESCs, whereas the E-proteins TCF3, 4 and 12, which are known to heterodimerize with TWIST1 (and many other bHLH TFs) to bind E-box motifs^40,47^, are expressed in both cell types, consistent with their broad expression across tissues (**Figure 2E**). Among the HD TFs, ALX1, ALX4, MSX1, and PRRX1 are most highly and selectively expressed in our hCNCCs, consistent with their closest resemblance to mesenchymal CNCCs of the anterior facial region^29^ (**Figure 2E**).

Accordingly, we created a panel of isogenic hESC lines with each TF endogenously and homozygously tagged with the dTAG-inducible FKBP12^F36V^ degron^48,49^, a V5 epitope tag, and in one case also the fluorophore mNeonGreen^50^, which we could then differentiate to hCNCCs *in vitro*^51^ (**Figure 3A**). This approach allows for acute or long-term depletion of each TF (**Figure 3B**) and—through the addition of a common V5 tag—for comparative studies of TF levels and DNA binding. Using CRISPR/Cas9 with adeno associated virus (AAV)-mediated delivery of homology-directed repair template^52^, we successfully tagged TWIST1, ALX1, MSX1, and PRRX1 and confirmed that we could induce near-complete depletion by adding dTAG^V^-1 molecule to the media (**Figure 3B**; **Figure S3A**). The aforementioned tagging strategy did not significantly disrupt baseline TF levels; for TWIST1 and MSX1, protein levels actually increased upon tagging (**Figure S3B**). Based on previous studies^42,53^, we suspected ALX1 and ALX4 might have overlapping functions, so we generated multiple independent clonal lines with nonsense mutations in *ALX4* on top of the *ALX1^FV^* tagged background, as we were unable to degron-tag ALX4 (**Figure 3C**; **Figure S3C**).

**Figure 3.**
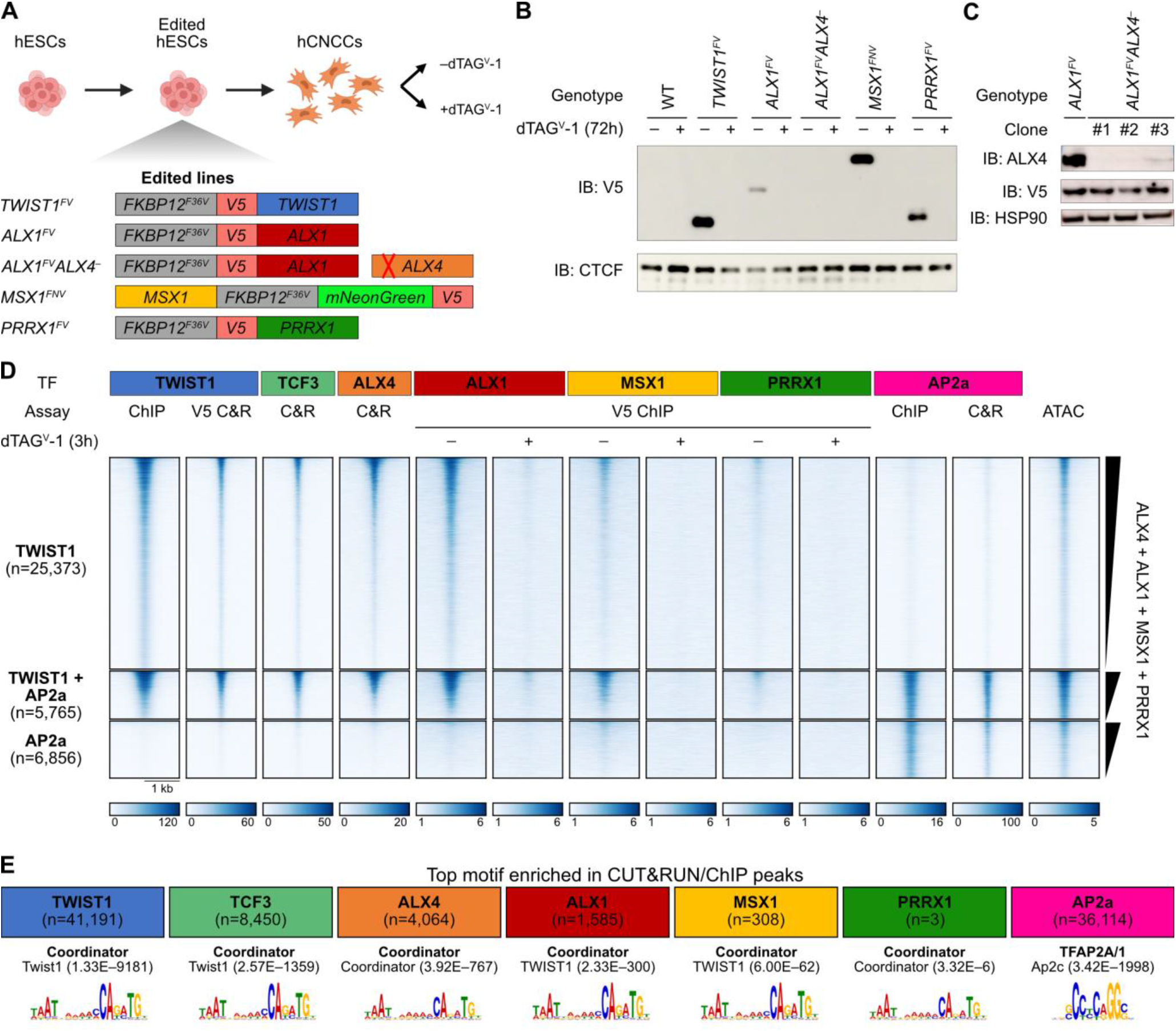
Multiple homeodomains co-bind Coordinator motif with TWIST1. **A**. Schematic of endogenous tagging TFs with the FKBP12^F36V^ degron and V5 epitope tag in human embryonic stem cells (hESC) followed by differentiation into cranial neural crest cells (hCNCC) and treatment with or without dTAG^V^-1. The ALX4 gene was instead knocked out by a frameshift mutation. **B**. Confirmation of TF tagging and depletion upon dTAG^V^-1 addition by Western blot for V5 epitope, with CTCF as a loading control. IB, immunoblot. **C**. Confirmation of ALX4 knockout in three independent clones by Western blot, with HSP90 as a loading control. **D**. Homeodomain TFs bind DNA at TWIST1 bound sites at variable occupancies. Heatmap shows promoter-distal binding sites for TWIST1 and/or AP2a. Assay indicates whether data shown is from ChIP, CUT&RUN (C&R), or ATAC, and whether an endogenous or V5 antibody was used. Rows are ranked by the sum of the homeodomain signals from undepleted cells. In the scale bar, units are reads per genome coverage, except for ATAC data, which is in signal per million reads. **E**. Homeodomain TFs bind the Coordinator motif. The top enriched motif (by AME) is shown for each TF, with p-values in parentheses.

Having developed these resources, we performed ChIP-seq and CUT&RUN to assess DNA binding profiles of these tagged TFs, plus ALX4, TCF3 (a heterodimerization partner of TWIST1), and the positive control AP2a (encoded by *TFAP2A*, a key neural crest TF^45^) using endogenous antibodies. We first used binding sites for TWIST1 and AP2a as reference points, grouping the distal regulatory regions into those bound by TWIST1 or AP2a only (**Figure 3D**, top and bottom heatmaps, respectively) or those co-bound by both (**Figure 3D**, middle heatmaps). As expected, binding of the TWIST1 heterodimerization partner TCF3 is correlated with that of TWIST1. For all four tested HD TFs, DNA binding at TWIST1 sites clearly exceeds that at AP2a-only sites despite comparable accessibility with TWIST1-only sites. The ChIP signal is minimal upon dTAG^V^-1 addition, underscoring the specificity of the V5 ChIPs. However, the strength of ChIP signal is reproducibly distinct between the tagged HD TFs, with strongest signal for ALX1, intermediate signal for MSX1, and only weak signal for PRRX1. We note that this ranking is not concordant with that of the TF protein levels by Western blot for the shared V5 epitope, as ALX1 has the lowest relative abundance but strongest binding (**Figure 3B**). ALX4 shows similar binding patterns as well, though we could not directly compare its chromatin occupancy level with that of other HDs, as in the absence of tags, we had to use an endogenous ALX4 antibody for the analysis.

As an orthogonal approach, we called peaks for each TF (using the depleted samples as negative controls when available) and searched for enrichment of known motifs (as before, clustering similar motifs) compared to shuffled controls (**Figure 3E**). The top motif cluster for TWIST1, TCF3, and all tested HD TFs is Coordinator, confirming that these HD TFs predominantly bind DNA with TWIST1. Together these data indicate that TWIST1 can bind Coordinator sites with multiple HD TFs including ALX4, ALX1, MSX1, and PRRX1, albeit at varying occupancies.

### TWIST1 facilitates homeodomain TF binding, chromatin opening, and enhancer activity

To investigate the mechanism and function of TF cooperation at Coordinator, we studied how depletion of each Coordinator-binding TF impacts chromatin states and the binding of other TFs. We first focused on TWIST1, given its central role as the key bHLH factor binding Coordinator across cellular contexts. We began with acute depletions ranging from 1 hour to 24 hours in hCNCCs, and performed ChIP-seq to measure TWIST1 binding, CUT&RUN for ALX4 binding (with an endogenous antibody), ATAC-seq to measure chromatin accessibility, and ChIP-seq for H3K27ac as a mark correlated with enhancer/promoter activity (**Figure 4A**).

**Figure 4.**
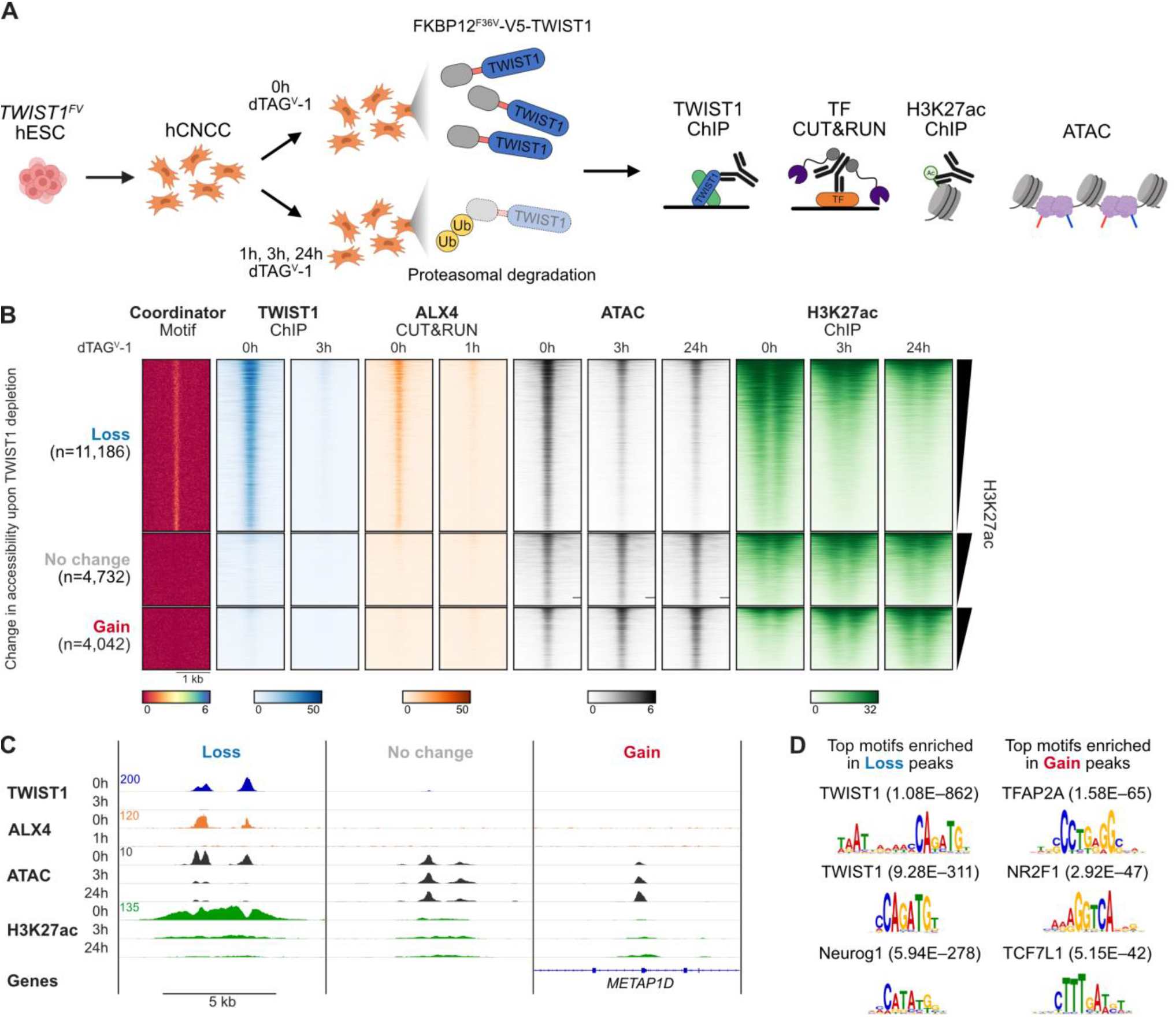
TWIST1 opens chromatin for homeodomain TFs and enhancer acetylation. **A**. Schematic of acute depletion experiments and follow-up assays. **B**. Heatmap of Coordinator motif enrichment, TF binding, chromatin accessibility (ATAC), and H3K27 acetylation (H3K27ac) at distal enhancers grouped by their change in accessibility upon TWIST1 depletion. In the scale bar, units are reads per genome coverage, except for the Coordinator motif, which is in log10 p-value, and ATAC data, which is in signal per million reads. One representative replicate of two independent differentiations is shown. **C**. Browser tracks of example enhancers with loss, no change, or gain of accessibility upon TWIST1 depletion. Coordinates in hg38: Loss, chr17:70,668,899-70,678,127; No change, chr11:44,958,683-44,968,011; Gain, chr2:172,058,768-172,068,096. **D**. Top three motif clusters enriched in enhancers with loss or gain of accessibility upon TWIST1 depletion compared to those with no change (by AME, analysis of motif enrichment) with their p-values.

TWIST1 depletion rapidly reshapes the chromatin landscape, with 36,290 regions losing accessibility and 17,054 regions gaining accessibility within 3 h (**Figure S4A**). The change in accessibility is mostly complete within 3 h (**Figure S4B**), so we combined the 3 h and 24 h differentially accessible peaks to define a set of sites with loss vs. gain of accessibility. Among candidate enhancers (promoter-distal peaks with robust H3K27ac signal), 11,186 sites lose accessibility, 4,042 sites gain accessibility, and 4,732 do not significantly change (**Figure 4B**, example browser tracks shown in **Figure 4C**). Regions losing accessibility are highly enriched for the Coordinator motif and TWIST1 binding, whereas those gaining accessibility lack TWIST1 binding and are most enriched for AP2a and NR2F1 motifs, suggesting these effects are indirect (**Figure 4B-D**). Changes in accessibility are correlated with changes in H3K27ac (**Figure 4B**, **Figure S4C**, r = 0.834 for 3 h, 0.896 for 24 h). In particular, loss of TWIST1 leads to a substantial depletion of H3K27ac within hours, consistent with an activating role of TWIST1 at *cis*-regulatory elements (**Figure S4D**). Furthermore, TWIST1 depletion eliminates enhancer reporter activity of a well-characterized *SOX9* enhancer dependent on the Coordinator motif^28^, confirming that TWIST1 is required for bona fide enhancer activity (**Figure S4E**). Importantly, TWIST1 depletion largely abrogates DNA binding of ALX4 at Coordinator sites in just 1 h (**Figure 4B,C**; **Figure S4D**). Therefore, both HD factor binding and open/active chromatin states of *cis*-regulatory elements are dependent on TWIST1, consistent with our original hypothesis based on evolutionary comparisons that the *trans*-regulatory protein or complex recognizing Coordinator may play a large role in ‘coordinating’ enhancer activity in CNCCs^20^.

### Homeodomain TFs cooperate with TWIST1 to open chromatin at Coordinator sites

We next asked how the depletion of HD TFs affects open chromatin landscapes and TWIST1 binding at Coordinator. Given that for ALX4 we were only able to generate a constitutive knockout, to obtain comparable data across all TF perturbations, we performed differentiations of *ALX4*^-^ hESCs along with the ALX1, MSX1, PRRX1, and TWIST1 degron-tagged hESCs, in which we treated cells with dTAG^V^-1 from the beginning of differentiations to mimic a knockout. We harvested these cells for ATAC-seq and RNA-seq at an early hCNCC stage, to minimize indirect effects. Even in these long-term depletions, many of the observed effects are likely directly caused by HD dysfunction in mesenchymal CNCCs, as most of the aforementioned HD TFs are only expressed in CNCCs following their specification and delamination^42,54^ (with an exception of MSX1, which is expressed in the neural plate border precursor to CNCCs^55^). Supporting this idea, accessibility effects of the long-term TWIST1 depletion are well correlated with the acute 24 h depletion (r = 0.664; **Figure S5A**).

Consistent with the range in strength of DNA binding among HDs (**Figure 3D**), ALX1 depletion results in significant changes in accessibility at 6,195 peaks (FDR < 0.05), while ALX4 knockout causes changes at 4,284 peaks, MSX1 at 1,410 peaks, and the weakest binder, PRRX1, at none (**Figure S5B**). In general, all HD TF depletions have much weaker effects on accessibility than TWIST1 depletion, likely due to functional redundancy among them. Indeed, changes upon ALX1 and ALX4 losses are well-correlated (r = 0.651) (**Figure 5A**). These are also correlated, albeit less well, with effects of MSX1 loss (r = 0.462) (**Figure S5C**). Next, by comparing the undepleted ALX1^FV^ samples (in which both ALX1 and ALX4 are present) to the depleted ALX1^FV^ ALX4^-^ samples (in which both are lost), we inferred the effect of combined ALX1 and ALX4 loss on the ATAC-seq changes at the corresponding set of genomic targets. This comparison allowed detection of differential accessibility at a greater number of peaks (8,577), but with similar or slightly larger effect sizes than individual ALX1 or ALX4 loss (**Figure 5B**).

**Figure 5.**
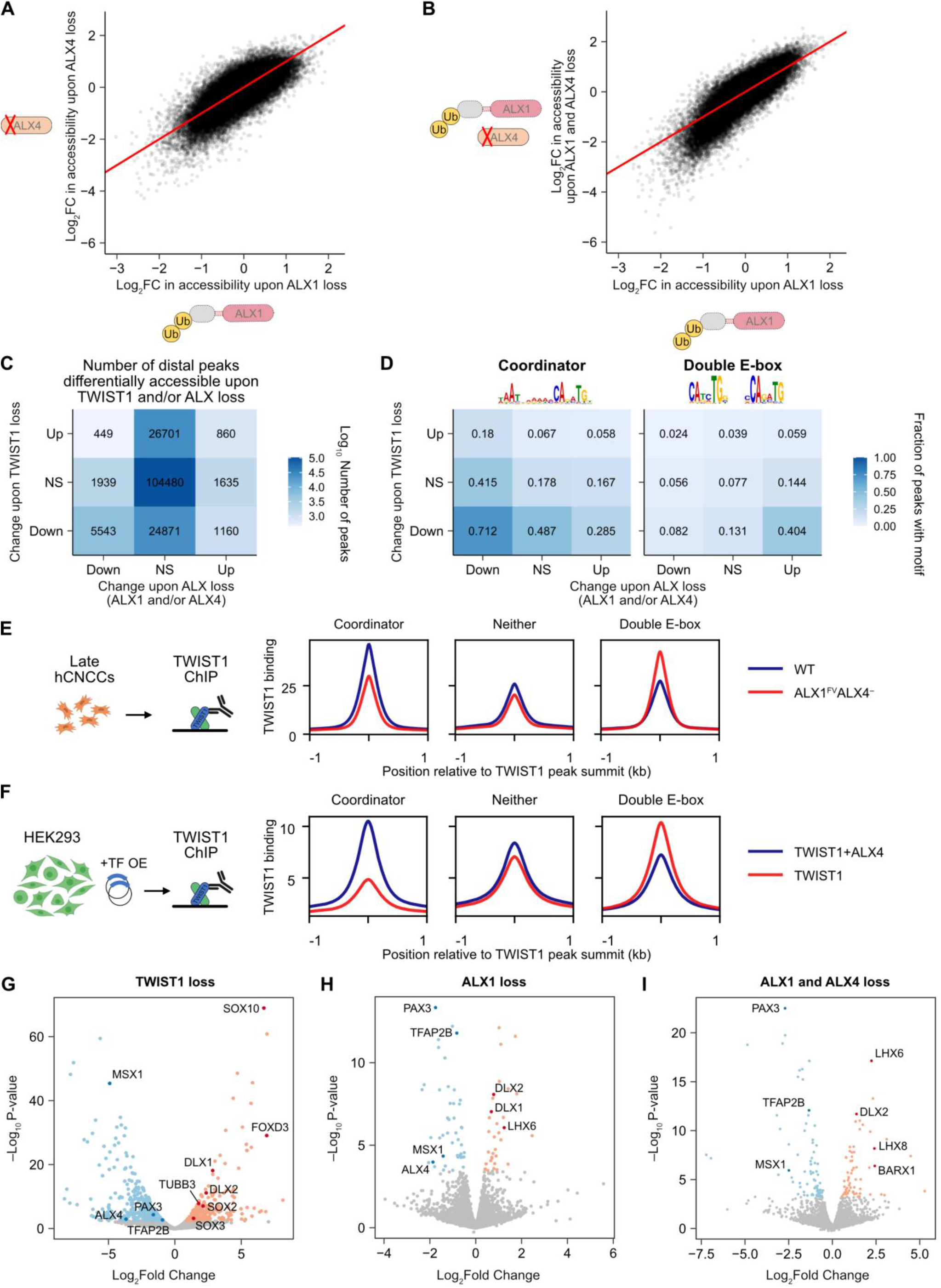
Homeodomain TFs stabilize TWIST1 binding at Coordinator sites. **A**. Correlation in Log_2_ fold change (FC) in accessibility upon loss of ALX1 (by long-term dTAG^V^-1 treatment) versus ALX4 (by knockout). All distal accessible regions with a significant change upon loss of any tested TF (ALX1, ALX4, MSX1, or TWIST1) are shown. Red line indicates y = x. **B**. Observed change in accessibility upon loss of both ALX1 and ALX4 is similar to the expectation based on the log sum of their individual effects. **C**. Most chromatin accessibility effects of ALX loss (ALX1 and/or ALX4) are concordant with (but are a subset of) those of TWIST1 loss. NS, not significant. **D**. Peaks with down regulation of accessibility upon TWIST1 and ALX loss are enriched in the Coordinator motif (left), while those that decrease upon TWIST1 loss but increase upon ALX loss are enriched in the double E-box motif (right). **E** and **F**. TWIST1 binding by ChIP-seq quantitatively shifts from Coordinator to double E box motif sites upon loss of ALX4 (without ALX1 depletion) in hCNCCs (**E**) or overexpression of TWIST1 alone rather than with ALX4 in HEK293 cells (**F**). **G** through **I**. Volcano plots of differential gene expression upon loss of TWIST1 (**G**), ALX1 (**H**), or ALX1 and ALX4 (**I**). ALX4 is excluded in (**I**). Upregulated genes are red and downregulated genes are in blue, with selected genes highlighted in darker colors.

sWe next asked how similar the effects of ALX loss on chromatin accessibility are to those of TWIST1 loss. Given the correlated effects of ALX1 and ALX4 loss (**Figure 5A,B**), we considered their combined effects, taking any ATAC-seq peak significantly affected by loss of ALX1, ALX4, or combined loss of both. As there are many more TWIST1-dependent peaks, most of these are not dependent on ALXs. However, of distal peaks downregulated upon ALX loss, the vast majority (5,543/7,931 = 70%) are concordant, or also downregulated upon TWIST1 loss, while few (449/7,931 = 5.7%) are discordant, or upregulated upon TWIST1 loss but downregulated upon ALX loss (**Figure 5C**). Distal peaks upregulated upon ALX loss lack this enrichment for concordance with TWIST1 effects, but these represent a minority (32%) of changes. The effects of MSX1 loss are also concordant with TWIST1 loss (**Figure S5E**). To find the DNA sequence features driving these concordant and discordant changes, we performed motif enrichment analyses on these different classes of accessible peaks. We found that the Coordinator motif is highly enriched in the TWIST1 and ALX-dependent peaks, underscoring that the main function of ALX1 and ALX4 in chromatin opening is indeed at Coordinator sites (**Figure 5D**). Of note, for the TFs that were degron-tagged, we also repeated the chromatin accessibility analysis upon acute loss of each TF, and observed minimal changes except for TWIST1 (**Figure S5D**) suggesting that, in contrast to TWIST1, the HD TFs are more important for establishing the chromatin landscape at Coordinator sites rather than for maintaining it after it has been established.

### Loss of homeodomain TFs titrates TWIST1 away from Coordinator towards the canonical double E-box sites

In addition to Coordinator, other motifs provide insight into the mechanisms underlying TWIST1-HD cooperation (**Figure 5D**; **Figure S5F**). In particular, the dominant feature of peaks that gain accessibility upon ALX loss but lose accessibility upon TWIST1 loss is the double E-box motif, which contains two E box motifs at a 5 bp spacing. The double E-box motif has previously been proposed to bind two copies of TWIST1:TCF3 heterodimers^40^ and we found it highly enriched in the top TWIST1 binding sites in the HD-negative RS4;11 cells (**Figure 5D**; **Figure S2E**). Thus, ALX loss appears to quantitatively redirect TWIST1 or its chromatin opening capacity away from Coordinator sites and towards double E-box sites.

To substantiate this observation and determine whether the distribution of TWIST1 binding at Coordinator vs double E-box sites is affected by the ALX loss, we performed TWIST1 ChIP-seq in *ALX1^FV^ ALX4^-^* hCNCCs (without ALX1 depletion) and compared the binding to that of WT cells (**Figure 5E**). TWIST1 binding signal is reduced at sites with the Coordinator motif but increases at sites with the double E-box motif. These changes are quantitative rather than qualitative, potentially due to the overlapping and partially redundant functions of HD TFs, which may retain some TWIST1 at Coordinator even in the absence of ALX4. To confirm this finding in a cellular context without redundancy, we overexpressed TWIST1 with or without ALX4 in HEK293 cells (which lack appreciable expression of TWIST1 or most HD TFs) and then performed TWIST1 ChIP-seq. As we saw in hCNCCs but to a greater extent in this overexpression context, TWIST1 binding to the Coordinator motif decreased in the absence of ALX4, whereas binding to the double E-box motif increased (**Figure 5F**).

### Shared transcriptional functions of TWIST1 and ALX factors

To assess the transcriptional functions of TWIST1 and HD factors in our *in vitro* hCNCC differentiation model, we used RNA-seq to identify genes significantly affected by the pertubation of TWIST1, ALX1, or both ALX1/4 (**Figure 5G-I**). Consistent with previous mouse studies^56,57^, the most significant effect of TWIST1 loss in our hCNCCs is an increase in *SOX10* expression (**Figure 5G**). SOX10 is an early CNCC regulator that becomes selectively downregulated as CNCC become biased toward mesenchymal rather than neuroglial lineages^57,58^. *FOXD3*, another early neural crest specifier gene^59,60^ whose expression decreases in post-migratory CNCCs^42,54^, is also highly upregulated upon loss of TWIST1, as are neural progenitor markers *SOX2/3* and the neuronal marker *TUBB3*, suggesting a defect in mesenchymal specification. Meanwhile, the loss of ALXs (which are expressed primarily in the anterior-most CNCC forming the frontonasal prominences) leads to upregulation of TF genes normally expressed only in more posterior parts of the face, such as *DLX1, DLX2, LHX6, LHX8*, and *BARX1*, and to downregulation of TF genes normally most abundant in the anterior-most regions of the face, such as *PAX3*, *TFAP2B*, and *ALX4* (the latter upon ALX1 depletion) (**Figure 5H,I**). This suggests that HD TFs associated with anterior facial identity (namely ALXs) promote expression of genes associated with this identity, while repressing expression of those associated with alternative, more posterior facial identities, as seen in a recent Alx1 null mouse^42^.

Notably, there is a substantial overlap between TWIST1 and ALX-responsive genes, with a subset of position-specific genes (*DLX1/2, PAX3, TFAP2B*) being regulated by TWIST1 as well as ALXs (**Figure 5G-I**). Furthermore, *MSX1*, a gene encoding HD TF broadly expressed throughout the face and limb buds and associated with mesenchymal cell identity^61^, is downregulated upon loss of ALXs as well as TWIST1. This overlap is representative of overall concordance between TWIST1 and ALX transcriptional changes: genes downregulated upon ALX loss are enriched for downregulation upon TWIST1 loss as well, in both long-term and acute depletions (**Figure S5G**). Note that MSX1 loss affects mesenchymal specification, with upregulation of neural progenitor markers *SOX2* and *SOX3* as seen with TWIST1 loss (**Figure S5H**), but generally has fewer effects than loss of ALXs, so shared activation of *MSX1* cannot explain most of the overlap in ALX and TWIST1 functions. These results suggest that TWIST1 and HD TFs co-binding at Coordinator sites drives shared transcriptional functions and may serve to integrate regulatory programs for lineage and regional identities during facial development.

### The Coordinator motif guides contact and cooperativity between TWIST1 and HD TFs

We next investigated biochemical and structural mechanisms underlying cooperative co-binding of TWIST1 and HD factors at Coordinator sites. We first considered cooperativity mediated by protein protein interaction between TWIST1 and HD proteins. We used immunoprecipitation-mass spectrometry (IP-MS) to systematically identify proteins that interact with TWIST1 in hCNCCs, using a chromatin extraction protocol that minimizes the extraction of DNA (**Figure 6A** and **Table S3**). Consistent with previously published results, we find that TWIST1 forms stable heterodimers with its E-protein partners TCF3, TCF4, and TCF12^40,47,62^. However, TWIST1 lacks interactions with ALXs or other HD TFs, and this was confirmed by the reciprocal IP-MS experiments pulling down the HD TFs (**Figure S6A** and **Table S3**).

**Figure 6.**
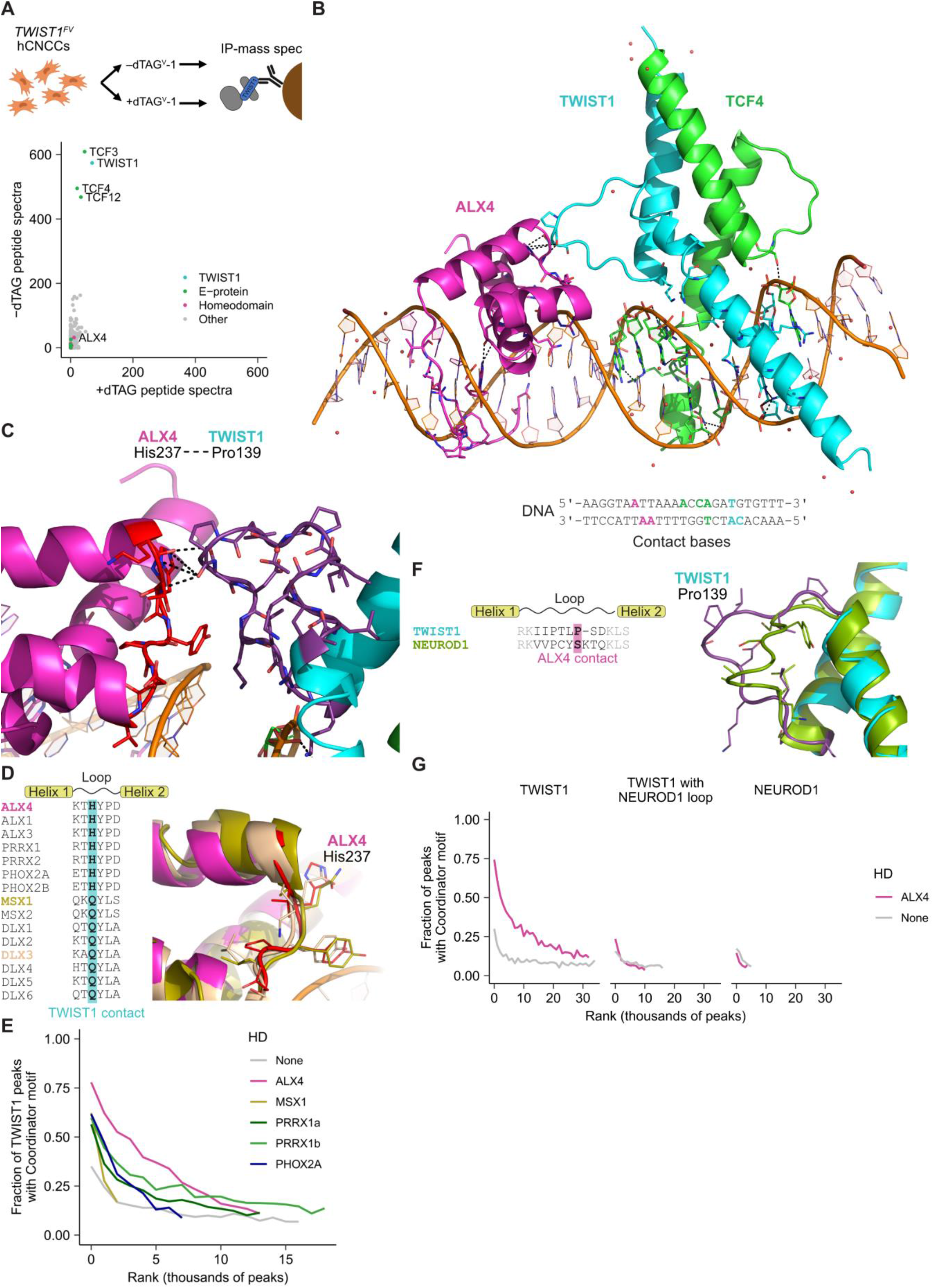
The Coordinator motif guides TWIST1-homeodomain contact and cooperativity. **A**. Immunoprecipitation-mass spectrometry (IP-MS) for TWIST1 using the V5 tag, in undepleted (-dTAG, y-axis) versus depleted (+dTAG, x-axis) hCNCC protein extracts. Plotted data are the sum of two biological replicates. **B**. 3D structure of TWIST1 (aa101-170), TCF4 (aa565-624), and ALX4 (aa210-277) bound to the Coordinator DNA sequence. DNA bases recognized by the TFs are highlighted in their respective colors: cyan for TWIST1, green for TCF4, and magenta for ALX4. **C**. Zoomed-in view of the contact between ALX4 residue His237 and TWIST1 residue Pro139. **D**. Sequence alignment of selected homeodomain TF loop sequences (left) with the TWIST1 contact residue highlighted in cyan, and structural alignment of ALX4 (magenta with red loop) with MSX1 (PDB: 1IG7) in gold, and DLX3 (PDB: 4XRS) in beige (right). **E**. Homeodomains shift TWIST1 toward Coordinator to varying degrees. Indicated Flag-tagged HD proteins were co-expressed with TWIST1 in HEK293 cells and TWIST1 binding was measured by ChIP-seq (see **Figure S6B** for protein levels). TWIST1 ChIP-seq peaks are ranked from strongest to weakest (left to right) and grouped into bins of 1000 peaks. **F**. Sequence alignment of TWIST1 and NEUROD1 loops (left), and structural alignment of TWIST1 (cyan with purple loop) with NEUROD1 (PDB: 2QL2) in green (right). **G**. TWIST1 loop is required for Coordinator binding. V5-tagged TWIST1, NEUROD1 or TWIST1 with NEUROD1 loop were expressed in HEK293 with or without ALX4 (see **Figure S6C** for protein levels). ChIP-seq peaks are ranked from strongest to weakest (left to right) and grouped into bins of 1000 peaks.

This observation suggested that cooperativity between TWIST1 and ALX proteins may be guided by the Coordinator motif DNA sequence. To explore this possibility, we solved an X-ray crystal structure of TWIST1, TCF4, and ALX4 DNA-binding domains co-bound to the consensus Coordinator motif, at 2.9Å resolution (**Figure 6B**). As expected, each TF is bound to its canonical motif, with a TWIST1-TCF4 heterodimer bound to the E-box and ALX4 bound to the HD monomer motif within Coordinator. Within the bHLH dimer, TWIST1 binds the side of the E-box motif further from the HD motif, allowing its loop to contact ALX4 (**Figure 6C**). The direct contact interface between the TWIST1 and ALX4 loops is one amino acid on each protein, with TWIST1 proline 139 interacting with ALX4 histidine 237. This lack of an extensive protein-protein binding interface is consistent with the absence of strong interactions in solution detectable by IP-MS, and further supports DNA-guided cooperativity between TWIST1 and ALX4.

The amino acid residues, and more broadly the loops, involved in the TWIST1-HD contact are not invariant in amino-acid sequence, even among TFs with highly similar DNA binding motifs (**Figure 6D,F**). To assess whether these variable loops form distinct structures, we aligned our TWIST1-TCF4-ALX4 Coordinator structure to previously solved (individual) HD and bHLH structures. Despite amino acid differences at the contact residue position (i.e. His to Gln substitution), MSX1 (PDB: 1IG7) and DLX3 (PDB: 4XRS) both form highly similar structures as ALX4 (**Figure 6D**). While the amino acid identity could impact the affinity of the contact, this result is consistent with our ChIP data suggesting that MSX1 and PRRX1 can also bind DNA at many of the same sites as ALX1/4 in hCNCCs, albeit at much reduced occupancies compared to the ALX factors (**Figure 3D**). To further test if these additional HD TFs can indeed direct TWIST1 binding towards Coordinator, we transfected plasmids encoding TWIST1 with one of ALX4, MSX1, PRRX1 (both major splice isoforms), or PHOX2A into HEK293 cells and performed TWIST1 ChIP-seq. All tested HD TFs are capable of increasing TWIST1 binding to the Coordinator motif, but none are as potent as ALX4 (**Figure 6E**), despite being expressed at comparable or higher protein levels (**Figure S6B**).

Among bHLH factors, neurogenic factors such as NEUROD1 stand out as having some of the most similar E-box DNA binding motifs to TWIST1 (**Figure S2A**). Yet, the NEUROD1 loop sequence and structure differ substantially from that of TWIST1 (**Figure 6F**). We reasoned that if, as our structure suggests, the loop contact plays a key role in Coordinator-guided cooperativity between bHLH and HD, then NEUROD1 itself or TWIST1 in which the ALX4-contacting loop has been replaced with the corresponding NEUROD1 loop may not be able to bind Coordinator in the presence of HD TFs. To test this, we transfected HEK293 cells with cDNAs encoding V5-tagged TWIST1, NEUROD1, or TWIST1 with a NEUROD1 loop, each with or without ALX4, ensuring that bHLH protein levels were comparable across samples (**Figure S6C**). We then performed ChIP-seq for the V5 tag. All three bHLH proteins are capable of binding chromatin at their known E-box motif (**Figure S6D,E**). Furthermore, as we have seen with an endogenous TWIST1 antibody, TWIST1 binds the Coordinator motif robustly only in the presence of ALX4. However, neither full length NEUROD1 nor TWIST1 with the NEUROD1 loop bind Coordinator appreciably even in the presence of ALX4 (**Figure 6G**). Collectively, these results illustrate how the Coordinator motif guides cooperative binding of TWIST1 and HD TFs, in a manner dependent on the features of the TWIST1 loop.

### The roles of Coordinator-binding TFs in facial shape variation

We initially identified Coordinator motif through the analysis of enhancer divergence between human and chimpanzee cranial neural crest (**Figure 1A**)^20^. As we have now uncovered the *trans*-regulatory complex that binds Coordinator, we aimed to assess the potential impacts of the identified TFs and their genomic targets on human phenotypic variation. Our previous genome-wide association study (GWAS) uncovered over 200 loci associated with normal-range variation in facial shape among individuals of European ancestry and revealed enrichment of face shape-associated genetic variants in CNCC enhancers^63^. To assess the contribution of Coordinator-binding TFs to human facial variation, we used two orthogonal approaches. In the first approach, we focused on enrichment of facial shape heritability at genomic targets regulated by Coordinator-binding TFs. In the second approach, we investigated the phenotypic impact of genetic variants at the loci encoding Coordinator-binding TFs themselves.

To assess contributions of specific sets of genomic regions responsive to TF losses, we used linkage disequilibrium score regression (S-LDSC) to determine the heritability enrichment of each set of regions compared to: (i) an accessibility-matched control set of hCNCC distal ATAC peaks (control) or (ii) the entire set of hCNCC distal ATAC peaks (all peaks) (**Figure 7A**). We first tested the set of distal regions differentially accessible within 3 h of acute TWIST1 depletion, separately assessing the up- and downregulated peaks. The downregulated peaks are 25.6-fold enriched over the genome, in contrast to 2.44-fold enrichment in the control peaks (p = 2.47E-6, downregulated vs matched control peaks, t-test) and 9.35-fold enrichment across all peaks (p = 6.63E-5, downregulated vs all peaks, t-test). In contrast, the upregulated peaks have a lower enrichment than either the matched or full control sets (1.78-fold vs 7.72-fold and 9.35-fold). We observed similar results for the peaks differentially accessible upon long term TWIST1 loss. Despite lower statistical power due to the smaller number of regions, we nevertheless detected significant enrichment in peaks downregulated upon ALX1+ALX4 loss compared to all hCNCC ATAC peaks (p = 0.0395, t-test). As a negative control, we analyzed the same genomic regions for enrichment of an unrelated trait, height. Height does not show the same pattern of enrichment in downregulated peaks even though height GWAS signal is enriched in hCNCC distal ATAC peaks overall, likely due to shared programs for skeletal development being involved in both traits (**Figure S7A**). These results indicate that genetic variation in the Coordinator-containing regions whose accessibility are regulated by TWIST1, ALX1, and ALX4 ultimately modulates human facial shape.

**Figure 7.**
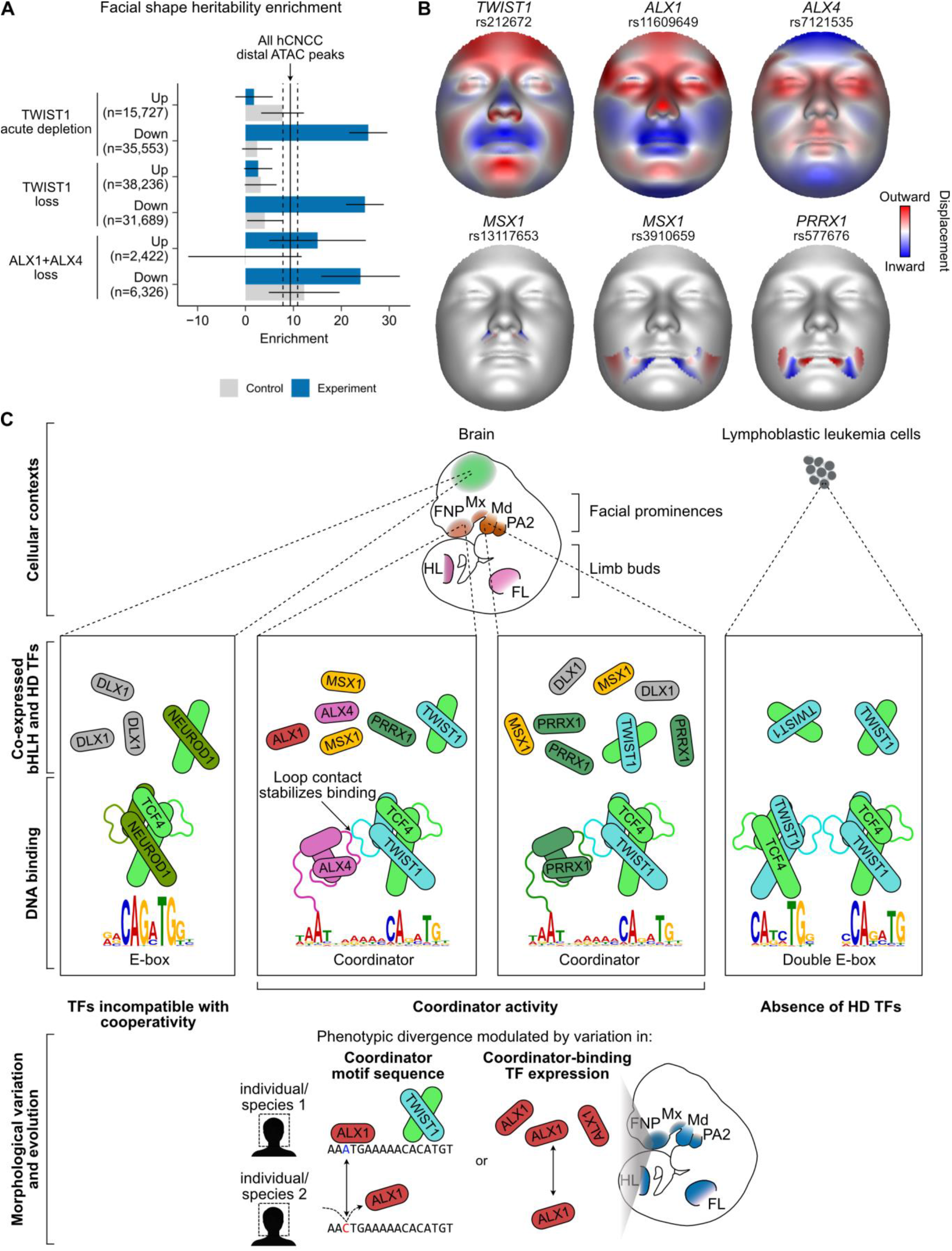
The roles of Coordinator-binding TFs in facial shape variation. **A**. Facial shape heritability enrichment at TWIST1- and ALX-dependent regulatory regions. Fold enrichment of facial shape-associated SNPs in distal ATAC peaks differentially accessible upon TF depletion or loss, with accessibility-matched control sets. Vertical line indicates the enrichment in all hCNCC distal ATAC peaks, with flanking dashed lines indicating error bars. Error bars represent s.e.m. **B**. Facial shape effects associated with genetic variants at loci encoding Coordinator-binding TFs. Facial shape effects of each lead SNP near Coordinator-binding TF genes, shown as normal displacement (displacement in the direction normal to the facial surface) for the facial region with highest significance for each SNP. Red colored areas shift outward and blue colored areas shift inward. **C**. Model of how DNA-guided cooperativity confers specificity among cellular contexts. The developing brain lacks TWIST1 and instead expresses neurogenic bHLH TFs like NEUROD1 which bind E-box motifs but cannot cooperate with co-expressed HDs. Different face and limb regions express TWIST1 with different HD partners capable of cooperativity at Coordinator motif. Lymphoblastic leukemia cells express TWIST1 but lack HD partners, so TWIST1 binds double E-box motif instead.

The analysis of facial shape GWAS signals revealed that loci encoding each of the Coordinator-binding TFs analyzed in this study (i.e. *TWIST1*, *ALX1*, *ALX4*, *MSX1*, and *PRRX1*) have at least one facial shape associated SNP in the non-coding region nearby, enabling us to examine the effect of modulating TF expression on human facial shape (**Figure 7B**; **Figure S7B**). The multivariate analytic approach of the facial shape GWAS allows for assessment of both global and local effects of genetic variants on facial shape by quantifying these effects and their significance over segments corresponding to distinct facial regions (**Figure S7B**)^63,64^. Facial effects can also be visualized by showing shape displacement associated with the minor vs major allele of a given genetic variant. When such analysis was performed for facial GWAS SNPs at Coordinator-binding TF loci, we observed that variants at the *TWIST1*, *ALX1*, and *ALX4* loci have broad and overlapping effects on facial shape, with the most significant effects spanning the entire face (**Figure 7B**; **Figure S7B**). In contrast, the SNPs at the *MSX1* and *PRRX1* loci have more localized effects near the mouth, which may be indicative of association with the maxilla shape. Our observations indicate that all loci encoding TF components of the Coordinator *trans*-regulatory complex are implicated in human facial variation. Furthermore, SNPs at the loci encoding TFs with the strongest regulatory input into Coordinator function (i.e. TWIST1 and ALX factors) show the most broad ranging facial effects. Together, these observations link Coordinator-binding TFs and their genomic targets to human phenotypic variation (**Figure 7C**).

## Discussion

Our work demonstrates how a composite TF motif called Coordinator guides selective co-binding by bHLH and HD factors (**Figure 7C**), thereby integrating established functions of these TFs in defining cell type^56,57,65^ (TWIST1) and positional^7,42,66^ (HDs) identities in the developing face and limb mesenchyme. Our study has important implications for evolution and variation of these mesenchymal structures and illustrates how DNA-guided cooperativity can facilitate increased regulatory specificity and selective functional interactions between TFs in the absence of stable protein complex formation in solution. Thus, the Coordinator motif and its *trans*-regulatory complex provide a novel paradigm for dissecting the mechanisms and implications of DNA-guided TF cooperativity in cellular contexts, which remains poorly understood beyond the extensively studied OCT4-SOX2 example.

### Coordinator motif and its binding factors in development, evolution, and disease

Although we first discovered the Coordinator motif through comparisons of human and chimpanzee CNCCs^20^, we now show that Coordinator is not restricted to primates nor the developing face. Instead, across cell types, Coordinator is selectively enriched at the *cis*-regulatory regions of the undifferentiated mesenchymal cells from both face and limb buds, which have distinct embryonic origins (neural crest vs mesoderm, respectively) but share expression of many key TFs. Across species, we detected Coordinator enrichment in mouse and chick limb bud mesenchyme (**Figure S1D**)^67^, suggesting an evolutionarily conserved role in vertebrates. Furthermore, the TFs that bind Coordinator (as well as their face and limb expression patterns) are also conserved among vertebrates^68–70^. Thus, the Coordinator motif appears to be one of the key unique *cis*-regulatory features of the vertebrate embryonic mesenchyme.

The TFs binding Coordinator have well-documented roles in the development of the face and limbs, as shown both in mouse models and revealed by human genetics. For example, mouse knockouts of *Twist1*^56,71^, *Alx1*^42^, and *Alx4* (in combination with mutations of *Alx1* or *Alx3*)^42,72^ all have strong craniofacial phenotypes that most profoundly manifest in the anterior facial regions, which interestingly, also coincides with the strongest enrichment of the Coordinator in the frontonasal prominence (FNP), as compared to other facial prominences. Similarly, Twist1^73^, Alx^74^, Msx^61,75^, and Prrx^76,77^ factors are involved in limb development in the mouse, where analogous to craniofacial regions, Alx4 marks the anterior-most region of the limb bud^78^, while Msx and Prrx factors are more widely expressed. In humans, mutations in *TWIST1* are associated with the Saethre-Chotzen and Sweeney-Cox syndromes, characterized by facial dysmorphisms, craniosynostosis, and limb malformations^79,80^, mutations in genes encoding ALX TFs cause frontonasal dysplasias^22,23,81,82^, and mutations in *PRRX1* are associated with agnathia-otocephaly complex (absence of mandible)^24^. Furthermore, our combined observations that: (i) genetic variants at the loci encoding Coordinator-binding TFs are associated with normal-range facial variation in humans, (ii) genomic targets regulated by these TFs are enriched for facial shape heritability, and (iii) Coordinator sequence mutations drive divergence in CNCC enhancer activity between human and chimp, altogether suggest that *cis*-regulatory mutations that affect Coordinator motif or expression level of its associated TFs play an important role in mediating inter- and intra-species phenotypic divergence in face shape. This role in phenotypic variation is likely not restricted to humans or primates. For example, genetic variants in the *ALX1* locus seem to have contributed to beak shape evolution in Darwin’s finches^83^, while a bat *PRRX1* enhancer contributed to its elongated forelimbs^84^.

### Coordinating cell type and positional identities

Embryonic development requires placement of the right cell types in the right places. Coordinator-guided cooperativity between TWIST1, a well-known regulator of mesenchymal lineage, and HDs, many of which have been implicated in establishing or maintaining positional identity (such as along anterior-posterior or proximal-distal axes), may serve to coordinate cell type and positional information in the embryonic mesenchyme. TWIST1 is broadly expressed across the undifferentiated mesenchyme of the face and limb buds, where it has been shown to promote mesenchymal identity^56,57,71^, and prevent premature chondrogenic or osteogenic differentiation^85,86^. The central role of TWIST1 in mesenchymal potential of the cranial neural crest is underscored by the recent observations that overexpression of TWIST1 is sufficient to endow trunk neural crest with the ability to form mesenchymal derivatives^58^, and to reprogram *Ciona* pigment cells into cells resembling the ectomesenchyme in vertebrates^87^. Beyond the face and limbs, TWIST1 functions in other processes associated with the acquisition of mesenchymal identity, such as during epithelial-to-mesenchymal transition (EMT) in metastasizing cancer cells^65^, and gastrulation and mesoderm development in *Drosophila*^88,89^. However, across these diverse biological contexts, TWIST1 has been shown to bind canonical solo and double E-box motifs^40,90^. Thus, TWIST1 performs distinct cellular and organismal functions, perhaps through a range of molecular mechanisms, with Coordinator-guided cooperativity with HD TFs enabling functions specific to face and limb development.

In contrast to the broad expression of TWIST1 across the developing mesenchyme, expression of HD TFs is more regionally restricted (**Figure 2D**; **Figure 7C**). Notably, most of the face is devoid of canonical HOX gene expression. Instead, ALX and DLX family HD TFs are expressed in position-restricted patterns and involved in development and patterning of the anterior and posterior facial structures, respectively^43,91^. ALX or DLX TF expression often coincides with that of MSX and PRRX TFs, which are more broadly transcribed throughout the developing face^91^. Our results indicate that multiple co-expressed HD TFs are capable of co-binding with TWIST1 at the Coordinator sites, albeit at varying occupancies, with the ALX TFs showing the strongest binding (**Figure 3D**, **Figure 6E**). Furthermore, the observation that Coordinator enrichment and TWIST1 binding at Coordinator sites are detectable in regulatory regions of mandibular prominence, which preferentially expresses DLX factors but lacks ALXs (**Figure 1D**, **2D**), combined with the structural similarity of the DLX3 homeodomain helix 1-helix 2 loop with that of ALX4 (**Figure 6D**), suggest that in the developing jaw mesenchyme, TWIST1 likely also cooperates with the DLX proteins. However, our data hint that the strength of Coordinator binding may contribute to the incipient divergence of facial regions, as the anterior-most FNP exhibits the highest Coordinator motif enrichment among TWIST1 binding sites, while the mandibular prominence exhibits reduced enrichment (**Figure 2B**). Together with our observation that ALXs have the strongest cooperation with TWIST1 (**Figure 6E**), this may explain the prior observation that a conditional knockout of TWIST1 in the neural crest leads to the most dramatic phenotype (a near-complete loss) in the upper face derived from the FNP and maxillary prominences, while the mandible is less affected^56^. Interestingly, loss of either TWIST1 or ALX TFs in our CNCC model results in the upregulation of *DLX1* and *DLX2* mRNAs (**Figure 5G-I**). This suggests that once positional identity of the facial region is established, it is reinforced by the cooperative function of TWIST1 and ALXs at the expense of the alternative, more posterior identity, mediated by the DLX factors.

### Specificity of DNA-guided cooperativity among bHLH and HD TFs

Cooperation at Coordinator is remarkably selective among cell types and TFs, akin to the OCT4-SOX2 motif defining pluripotent stem cells. Even TFs with highly similar individual TF motifs that are co expressed with some of the same candidate partner TFs are unable to cooperate: NEUROD1 cannot cooperate with ALX4 (**Figure 6G**), and in the developing forebrain, the abundant DLX factors do not bind Coordinator despite nearby enrichment of neurogenic bHLH TF motifs (**Table S1**)^92^. Nevertheless, *in vitro*, other bHLH-HD TF pairs can co-bind composite motifs by CAP-SELEX^14^, so while Coordinator itself has not been seen in other cellular contexts, other TF pairs may be capable of co-binding distinct composite motifs with heretofore unexplored biological functions.

Cooperative potential is quantitative rather than binary—while many HDs can direct TWIST1 to Coordinator, they do so at lower efficiency than ALX4 (**Figure 6E**). This may explain why even though neuroblastomas exhibit high expression of PHOX2A/B, which can direct TWIST1 to Coordinator (**Figure 6E**), TWIST1 does not bind Coordinator very robustly in neuroblastomas (**Figure 2B**). ALXs appear particularly dependent on TWIST1 for their chromatin occupancy, with binding sites overwhelmingly enriched for Coordinator motifs rather than independent HD monomer or dimer motifs (**Figure 3E**), but other TFs may bind mostly at independent sites. Whether a given pair of TFs will preferentially bind at composite, cooperative sites versus their canonical monomeric or dimeric sites may depend not only on the strength of co-binding between the two partners, but also on the milieu of other TFs capable of functional (direct or indirect) interactions with one (or both) of the cooperating TFs. For example, for bHLH and HD these functional interactions may include their heterodimerization partners, such as E-proteins and TALE-type HD TFs^93^, respectively.

We find through structure and mutagenesis studies that a fairly weak contact between flexible loops can stabilize cooperative binding by TWIST1 and ALX4 at Coordinator sites (**Figure 6**, **Figure 7C**). The weak interaction between the loops is guided by the DNA, as we do not detect association between TWIST1 and ALX TFs in solution, even though robust binding between TWIST1 and its E-protein partners is seen under the same conditions (**Figure 6A**). The weakness and flexibility of the inter-loop contact may make it more easily modulated during evolution, and indeed the loop residues are variable among closely related bHLH and HD TFs. Replacing the loop in TWIST1 bHLH domain with that of NEUROD1 renders it incapable of cooperating with ALX4 (**Figure 6G**), without affecting ability of the bHLH domain to bind at canonical E-box sites (**Figure S6E**), underscoring the importance of the loop in mediating specificity of the cooperative binding. Interestingly, a recent study showed that the bHLH loop of the yeast pioneer factor Cbf1 is involved in nucleosome contacts critical for nucleosome binding^94^, suggesting that the bHLH loop may have pleiotropic biochemical functions. More work is needed to map the network of potential and observed cooperativity among TFs, but we propose that loops offer a mechanism that balances specificity with flexibility.

## Acknowledgments

We thank Katherine Xue, Raquel Fueyo, Christina Jensen, Tiffany Chern, and Liang-Fu Chen for critical feedback, Hannah Long for generating the pGL3-noSV40-humanEC1.45_min1-2_4xEboxMutant plasmid, and Seppe Goovaerts for assistance retrieving facial GWAS data. Mass spectrometry data were collected by the Vincent Coates Foundation Mass Spectrometry Laboratory, Stanford University Mass Spectrometry (RRID:SCR_017801). This work was supported in part by NIH P30 CA124435 utilizing the Stanford Cancer Institute Proteomics/Mass Spectrometry Shared Resource. Protein production was performed by the Protein Sciences Facility in the Karolinska Institutet Department of Medical Biochemistry and Biophysics.

pAAV-GFP was a gift from John Gray (Addgene plasmid # 32395)^95^, pDGM6 was a gift from David Russell (Addgene plasmid # 110660)^96^, PB-iNEUROD1_P2A_GFP_Puro was a gift from Prashant Mali (Addgene plasmid # 168803)^97^, and pCAG-NLS-HA-Bxb1 was a gift from Pawel Pelczar (Addgene plasmid # 51271)^98^.

This work was supported by HHMI-Damon Runyon Cancer Research Foundation Fellowship (DRG-2420-21), Stanford School of Medicine Dean’s Postdoctoral Fellowship, and NIH training grant 2T32AR007422-36A1 to S.K., Helen Hay Whitney Fellowship to S.N., Distinguished Professor Award from the Swedish Research Council to J.T., and NIH grant R35 GM131757, the Nomis Foundation, funding from the Howard Hughes Medical Institute, a Lorry Lokey endowed professorship, and a Stinehart Reed award to J.W.

## Author contributions

Conceptualization, S.K., J.T., and J.W.; Methodology, S.K., E.M., S.N., M.K.; Formal Analysis, S.K., E.M., and S.N.; Investigation, S.K., E.M., S.N., M.B., M.K., A.Popov, C.L., A.Pogson., J.T., and J.W.; Writing – Original Draft, S.K. and J.W.; Writing – Review & Editing, S.K., E.M., S.N., J.T., and J.W.; Visualization – S.K. and E.M.; Project Administration, J.W.; Supervision, J.W., J.T., and P.C.; Funding Acquisition, J.W., J.T., S.K., and S.N.

## Materials and Methods

### Lead contact

Further information and requests for resources and reagents should be directed to and will be fulfilled by the Lead Contact, Joanna Wysocka.

### Materials availability

Plasmids generated in this study will be deposited in Addgene upon peer-reviewed publication. All other reagents are available upon request.

### Data and code availability

All sequencing datasets have been deposited in NCBI GEO at accession GSE230319. Accession numbers of analyzed publicly available datasets are listed in **Table S2**. ENCODE datasets were downloaded from https://www.encodeproject.org/. CCLE data were downloaded from https://depmap.org/portal/download/all/, Release 22Q1 “CCLE_expression.csv” and “sample_info.csv”. Mass spectrometry peptide spectrum match counts are provided in **Table S3**. The TWIST1-TCF4-ALX4 crystal structure atomic coordinates and diffraction data have been deposited to Protein Data Bank under accession 8OSB. All original code have been deposited to Zenodo at https://doi.org/10.5281/zenodo.7847853.

### Cell culture

H9 cells (WiCell, WA09, RRID:CVCL_9773) were cultured in feeder-free conditions, in mTeSR1 medium (Stem Cell Technologies, 85850) on Matrigel Growth Factor Reduced (GFR) Basement Membrane Matrix (Corning, 356231) and passaged using ReLeSR (Stem Cell Technologies, 05872) every 4-6 days. Cells were switched to mTeSR Plus medium (Stem Cell Technologies, 100-0276) prior to and during genome editing and clonal expansion, but switched back to mTeSR1 before differentiation to CNCC. hESCs were differentiated to CNCC as previously described^20,28^. Briefly, hESC colonies were partially detached from the plate with collagenase IV (Gibco, 17104019) in Knockout DMEM medium (Gibco, 10829018) for 30-60 min and scraped to break up large colonies, and then cultured in Neural Crest Differentiation Medium (50%-50% v/v mixture of DMEM/F12 1:1 medium with L-glutamine, without HEPES (Cytiva, SH30271.FS) and Neurobasal medium (Gibco, 21103049) with 0.5x N2 NeuroPlex (Gemini Bio, 400-163) and Gem21 NeuroPlex (Gemini Bio, 400-160) supplements and GlutaMAX (Gibco, 35050061), and 1x antibiotic/antimycotic, and 20 ng/ml EGF (Peprotech, AF-100-15), 20 ng/ml bFGF (Peprotech, 100-18B), and 5 ug/ml bovine insulin (Gemini Bio, 700-112P)) for 11 days in bacterial-grade petri dishes, changing the plate to prevent attachment for 4 days and then leaving the cells unfed for two days to allow attachment, and then fed as needed at least every other day. At day 11, cells (now called ‘early hCNCC’) were harvested by treatment with Accutase (Sigma-Aldrich, A6964-100ML), strained to remove residual neuroectodermal spheres, and plated onto plates coated with 7.5 ug/ml human fibronectin (Millipore, FC010-10MG) and cultured in Neural Crest Maintenance Medium (Neural Crest Differentiation Medium with bovine insulin replaced by 1 mg/ml BSA (Gemini Bio, 700-104P)). These hCNCC were then passaged every 2-3 days upon reaching confluency, with cells in the third or subsequent passages defined as ‘late hCNCC’ and cultured with added 50 pg/ml BMP2 (Peprotech, 120-02) and 3 uM CHIR 99021 (Selleck, S2924).

dTAG^V^-1 (Tocris, 6914/5) was dissolved in DMSO at 5 mM and then diluted to 250 uM in 60% DMSO/40% water (v/v) before dilution to 500 nM for acute depletions (up to 1 day) or diluted directly from the 5 mM stock for long-term depletions. For acute depletion time courses, an equivalent amount of DMSO (0.12% v/v final) was added to all samples starting 24 h before harvest, and cells for all time points were harvested simultaneously.

RS4;11 cells (ATCC, CRL-1873, RRID:CVCL_0093) were cultured in RPMI-1640 medium (Gibco, 11875093) supplemented with 10% v/v FBS and 1x antibiotic/antimycotic.

HEK293 cells (ATCC, CRL-1573, RRID:CVCL_0045) and 293FT cells (Invitrogen, R70007, RRID:CVCL_6911) were cultured in DMEM high glucose medium with sodium pyruvate and L-glutamine, supplemented with 10% v/v FBS and 1x GlutaMAX, non-essential amino acids, and antibiotic/antimycotic.

O9-1 cells (Millipore, SCC049, RRID:CVCL_GS42) used for spike-in controls for ChIPs of TWIST1 depletions were cultured in Complete ES Cell Medium with 15% FBS (Millipore, ES-101-B), 25 ng/ml bFGF, and mLIF (Millipore, ESG1107).

### Oligonucleotides

Primers used in this study are listed in **Table S4**.

### Plasmids and cloning

AAV donor templates were cloned into the pAAV-GFP (Addgene plasmid # 32395) backbone by digesting pAAV-GFP with SpeI-HF (New England Biolabs, R3133S) and XbaI (New England Biolabs, R0145S) and performing Gibson assembly (New England Biolabs, E2611S) with PCR products of the ∼1 kb homology arms and tags. Flexible linkers (glycine-serine or glycine-alanine) of 5-11 aa were added in between the degron and epitope tags and the TF of interest.

Plasmids in the pCAG backbone used to overexpress TWIST1 and ALX4 in HEK293 cells were cloned by digesting the pCAG-NLS-HA-Bxb1 plasmid (Addgene plasmid # 51271) prepared from dam-/dcm- *E. coli* (New England Biolabs, C2925H) with SalI-HF (New England Biolabs, R3138S) and BclI (New England Biolabs, R0160S) followed by Gibson assembly with PCR products of desired inserts.

Plasmids in the pcDNA3.1 backbone used to overexpress V5-tagged TWIST1/NEUROD1 and ALX4 in HEK293 cells were cloned by PCR of the pcDNA3.1 backbone and desired inserts followed by Gibson assembly.

Coding sequences of MSX1 (NM_002448.3, OHu18516D), PRRX1a (NM_006902.5, OHu23742D), PRRX1b (NM_022716.4, OHu15551D), PHOX2A (NM_005169.4, OHu18020D) were ordered from Genscript. TWIST1 was amplified from H9 gDNA, with tags added following the second ATG at the beginning of the coding sequence. NEUROD1 was amplified from PB-iNEUROD1_P2A_GFP_Puro (Addgene plasmid # 168803). FKBP12^F36V^-V5 (for N-terminal tagging) was synthesized by Integrated DNA Technologies. FKBP12^F36V^-mNeonGreen-V5 (for C-terminal tagging) was amplified from pAAV-hSOX9-dTAG-mNeonGreen-V5 (Addgene plasmid #194971).

The pGL3-noSV40-humanEC1.45_min1-2_4xEboxMutant plasmid was generated by mutating all four E box motifs within Coordinator motifs *in silico* at the positions with greatest information content in the PWM. The sequence containing mutant EC1.45 E-box motifs was ordered from Twist Bioscience and cloned into the pGL3 luciferase reporter vector.

### AAV preparation

AAV production was performed by transfecting 293FT cells with 22 ug of pDGM6 helper plasmid (Addgene plasmid # 110660), 6 ug of donor template plasmid, and 120 ug polyethylenimine (Sigma Aldrich, 408719) diluted in Opti-MEM (Gibco, 31985070) in 1 ml total volume per 15-cm plate (4 plates were used per construct). Twenty-four hours after transfection, media was changed to media with 2% FBS. Three days after transfection, cells were harvested by scraping, triturated by pipetting up and down, centrifuged at 1000g for 20 min at 4°C, resuspended in 1.5 ml AAV lysis buffer (2 mM MgCl2, 10 mM NaCl) per 2×15 cm plates, and then flash frozen for storage. Samples were passaged through a 23 gauge needle and then freeze-thawed three additional cycles to lyse cells. Lysates were then treated with Benzonase (Millipore, 71205-3) for 1 h at 37°C with intermittent mixing, centrifuged at 2000g for 20 min at 4°C, and then the supernatant was flash frozen for storage at -80°C. OptiSeal tubes (Beckman Coulter, 362183) were filled from the bottom (with a blunt 18-gauge needle attached to a syringe), in order, with layers of 9.7 ml of 25% OptiPrep Density Gradient medium (Sigma-Aldrich, D1556-250ML) in 100 mM Tris pH 7.6, 1.5 M NaCl, 100 mM MgCl_2_; 6.4 ml of 41.7% OptiPrep in 100 mM Tris pH 7.6, 0.5 M NaCl, 100 mM MgCl_2_, and 12 ug/ml Phenol Red; 5.4 ml of 66.7% in 100 mM Tris pH 7.6, 0.5 M NaCl, 100 mM MgCl_2_, and 5.4 ml of 96.7% OptiPrep Density Gradient medium (Sigma-Aldrich, D1556-250ML) in 33.3 mM Tris pH 7.6, 167 mM NaCl, 33 mM MgCl_2_ with 0.012 mg/ml Phenol Red. Lysate was gently added on top, the tubes were filled with AAV lysis buffer, and centrifuged at 48,000 rpm at 18°C in a Beckman VTi 50 rotor for 70 min with max acceleration and braking at a setting of 9. The viral fraction above the 66.7%-96.7% OptiPrep interface was collected using an 18-gauge needle and syringe and then washed with cold PBS using an Amicon Ultra-15 100K filter (Millipore, UFC910008). Purified AAV was then flash frozen in aliquots for storage at -80°C. To calculate AAV titers, an aliquot was digested with Turbo DNase (Invitrogen, AM2238) per manufacturer’s instructions, inactivated with 1 mM EDTA final concentration and incubation at 75°C for 10 min, and then digested with proteinase K in 1 M NaCl and 1% w/v N-lauroylsarcosine at 50°C for 2h. Samples were then boiled for 10 min, and diluted in H2O to 1:20,000 and 1:200,000. DNA standards comprising 10^10^ - 10^3^ molecules were prepared using AAV6 backbone plasmids containing inverted terminal repeats. Quantitative PCR was carried out on standards and test samples using the LightCycler 480 Probes Master kit (Roche, 04707494001) with inverted terminal repeat probe and primer sequences indicated in **Table S4**.

### Genome editing

H9 cells were treated with 10 uM Y-27632 (Stem Cell Technologies, 72304) for at least 2 h prior to nucleofection, and then harvested as single cells with Accutase. For each editing experiment, 800,000 cells were nucleofected with 1.7 ul (17 ug) SpCas9 HiFi (Integrated DNA Technologies) and 3.3 ul of 100 uM annealed crRNA XT and tracrRNA (pre-incubated for 15 min at room temperature to form RNPs) and for generating ALX4 knockout, 2 ul of 100 uM ssDNA homology-directed repair (HDR) template, using the P3 Primary Cell 4D-Nucleofector X Kit L (Lonza, V4XP-3034) and the CA-137 program. When AAV was used to deliver HDR template, the AAV was diluted to 25,000 viral genomes per cell in medium and added to the plate before adding the nucleofected cells. Media was changed 4 h after nucleofection, and then cells were cultured until nearing confluency, at which point cells were diluted to single cells and plated at low densities (∼500 cells per well of a 6-well plate). Resulting colonies were picked and a portion of the cells lysed by QuickExtract (Lucigen, QE9050) and used to genotype by PCR with primers on either side of the insertion site (in most cases with one primer outside the homology arms; see **Table S4** for primer sequences) and gel electrophoresis or Sanger sequencing. Putatively edited colonies were confirmed by genomic DNA extraction using the Quick-DNA mini prep kit (Zymo, D3024) and Sanger sequencing. All gRNA and primer sequences are listed in **Table S4**.

### Transfection

HEK293 cells were transfected with Lipofectamine 2000 (Invitrogen, 11668019) at a ratio of 2.8 ul lipofectamine per ug of DNA, diluted in Opti-MEM. Cells were transfected with 2.5 ug DNA per well of a 6-well plate or 15 ug DNA per 10-cm plate 1-2 days after seeding, when they reached 70-90% confluency. Media was replaced 4-6h after transfection, and then cells were harvested for Western blot or chromatin immunoprecipitation at 24 h after transfection.

hCNCCs were transfected with FuGENE 6 (Promega, E2691) immediately after passaging, using 1 ul of FuGENE 6 per 3 ug of DNA and 100 ng DNA diluted in 50 ul Opti-MEM per well of a 24-well plate.

### Luciferase assay

hCNCCs were transfected with 0.5 ng pRL renilla control plasmid, 10 ng modified pGL3 reporter plasmid, and 89.5 ng carrier plasmid (pUC19) per well of a 24-well plate, in triplicate. Cells were lysed 24 h after transfection and assayed with the Dual-Luciferase Reporter assay kit (Promega, E1960).

### Western blot

Cells were washed with cold PBS, lysed by incubation for 10 min on ice in RIPA buffer (50 mM Tris pH 7.6, 150 mM NaCl, 1% Igepal CA-630, 0.5% sodium deoxycholate, 0.1% SDS) with 1x cOmplete EDTA free protease inhibitor cocktail (Roche, 11873580001), and sonicated for 6 cycles of 30s ON/30s OFF on high power using the Bioruptor Plus (Diagenode). Insoluble material was removed by centrifugation at >16,000g for 10 min at 4°C. The supernatant was quantified by BCA protein assay (Thermo, 23225) and then denatured by addition of 1x NuPAGE LDS Sample Buffer (Invitrogen, NP0007) and 100 mM DTT and heating to 95°C for 7 min. Samples were normalized by BCA quantifications and then loaded in 4-12% or 4-20% Novex Tris-glycine gels (Invitrogen) and run at 165V for ∼1 h in Tris-glycine buffer (25 mM Tris and 192 mM glycine) with 0.1% SDS. Gels were transferred onto nitrocellulose membranes (GE Healthcare) for 1 h at 400 mA in Tris-glycine buffer with 20% methanol, stained with 0.1% Ponceau S in 3% trichloroacetic acid, then blocked with 5% milk and 1% BSA in PBS with 0.1% Tween-20 (PBST) for 15 min at room temperature, and then incubated with primary antibody overnight at 4°C followed by horseradish peroxidase (HRP)-conjugated secondary antibody incubation for 1 h at room temperature, with 4 washes of PBST after each antibody incubation. Antibodies used include TWIST1 (Abcam, ab50887, RRID:AB_883294, 1:500), ALX4 (Novus Bio, NBP2-45490, 1:1000), ALX1 (Novus Bio, NBP1-88189, 1:1000), MSX1 (Origene, TA590129, 1:5000), PRRX1 (Santa Cruz Biotechnology, sc-293386, 1:500), CTCF (Cell Signaling, 2899, 1:2000), HSP90 (Cell Signaling, 4877, 1:2000), V5 (Abcam, ab206566, 1:2000), Flag (Sigma, F1804, 1:2000), HA (Abcam, ab9110, 1:2000), Donkey anti-Rabbit IgG (H+L) HRP (Jackson Immunoresearch, 711-035-152, RRID:AB_10015282, 1:3000), Goat anti-Mouse IgG (H+L) HRP (Jackson Immunoresearch, 115-005-003, RRID:AB_2338447, 1:3000). Chemiluminescence was performed with Amersham enhanced chemiluminescence (ECL) Prime reagent (Cytiva, RPN2232) and imaged with an Amersham Imager 800 (Amersham).

### ATAC-seq

Omni-ATAC was performed essentially as published^100^, with 30 min treatment with 200 U/ml DNase I (Worthington, LS006331) at 37°C prior to harvesting cells, using Ampure XP (Beckman Coulter, A63881) beads to clean up the DNA. Briefly, treated cells were harvested by Accutase, counted using the Countess II (Invitrogen), and 50,000 cells were collected by centrifugation at 500g for 5 min at 4°C. Cells were resuspended in lysis buffer (resuspension buffer (RSB), or 10 mM Tris-HCl pH 7.4, 10 mM NaCl, and 3 mM MgCl2, with 0.1% Igepal CA-630, 0.1% Tween-20, and 0.01% digitonin) for 3 min, then quenched by dilution with RSB with 0.1% Tween-20. Lysate was centrifuged for 10 min at 500g at 4°C and then resuspended in transposition buffer (25 ul TD buffer and 2.5 ul TD enzyme (Illumina, 20034197), 16.5 ul PBS, 0.01% digitonin, 0.1% Tween-20, and water up to 50 ul). Transposition reactions were performed for 30 min at 37°C, and then cleaned up with the DNA Clean & Concentrator-5 kit (Zymo, D4013) and eluted in 21 ul 10 mM Tris-HCl pH 8. DNA was then pre-amplified 5 cycles with NEBNext Ultra II Q5 master mix (New England Biolabs, M0544) with a cycling protocol of 72°C for 5 min, 98°C for 30s, and 5 cycles of 98°C for 10s, 63°C for 30s, 72°C for 1 min. Then 5 ul of the 50 ul reaction was used to run a qPCR reaction (with the same cycling protocol except the initial 72°C incubation) to determine the optimal number of PCR cycles for each sample. The remaining portion of the reaction was then amplified the appropriate number of cycles, and then subjected to two rounds of double-sided Ampure XP bead cleanup, with 0.5x/1.3x and 0.5x/1.0x bead ratios (numbers indicate bead ratios added in first and second steps). Libraries were quantified by Qubit dsDNA high sensitivity assay (Invitrogen, Q33231), run on a 5% polyacrylamide TBE gel to check the size distribution, and then pooled for sequencing.

### Chromatin immunoprecipitation

Cells (about 1 confluent 10-cm plate or ∼10-20 million cells) were crosslinked with 1% methanol-free formaldehyde (Pierce, 28908) in PBS for 10 min at room temperature and then quenched by adding 2.5 M glycine to 125 mM final concentration and incubating for 10 min. Cells were washed in PBS with 0.001% v/v Triton X-100, harvested by scraping, and collected by centrifugation for 5 min at 4°C. Cells were washed with PBS and flash frozen for storage at -80°C. Cell pellets were later thawed on ice for 30 min, and then sequentially resuspended in lysis buffer 1 (50 mM HEPES-KOH pH 7.5, 140 mM NaCl, 1 mM EDTA, 10% glycerol, 0.5% Igepal CA-630, 0.25% Triton X-100, 1x cOmplete EDTA-free protease inhibitor cocktail (PIC), 1 mM PMSF), lysis buffer 2 (10 mM Tris-HCl pH 8, 200 mM NaCl, 1 mM EDTA, 0.5 mM EGTA, 1x cOmplete EDTA-free protease inhibitor cocktail, 1 mM PMSF), and lysis buffer 3 (10 mM Tris HCl pH 8, 100 mM NaCl, 1 mM EDTA, 0.5 mM EGTA, 0.1% sodium deoxycholate, 0.5% N lauroylsarcosine, 1x PIC, 1 mM PMSF), with 10 min incubations in each buffer, with rotation. Lysates were sonicated for 10-15 cycles of 30s ON/30s OFF on high power using the Bioruptor Plus (Diagenode), then diluted in additional lysis buffer 3 and clarified by centrifugation for 10 min at max speed at 4°C. Triton X-100 was added to 1%, and a small aliquot was used to extract DNA to check chromatin yield and size distribution, by dilution in elution buffer (1% w/v SDS and 100 mM NaHCO3) and incubation with 200 mM NaCl and RNase A (Thermo, EN0531) at 65°C for 1 h, then proteinase K (Thermo, EO0491) at 65°C for 1 h, and clean up with the ChIP DNA Clean & Concentrator-5 kit (Zymo, D5205). DNA was quantified by Qubit dsDNA high sensitivity kit, and the remaining chromatin was then normalized for immunoprecipitations. For TWIST1 acute depletions, chromatin from O9-1 mouse CNCCs were added prior to ChIP at ∼10% of the total chromatin as a spike-in control. Antibodies used include TWIST1 (Abcam, ab50887), V5 (Abcam, ab15828), H3K27ac (Active Motif, 39133), Flag (Sigma-Aldrich, F1804), AP2a (Cell Signaling, 3215), AP2a (Novus Bio, NB100-74359). For H3K27ac, 5 ug of antibody was used per ChIP; for TFs, 9 ug of antibody was used per ChIP, except for dissected mouse embryos where 4.5 ug was used in half of the total ChIP volume. ChIPs were incubated overnight, then incubated for 4-6h with 100 ul Dynabeads Protein A (Invitrogen, 10002D) or Protein G (Invitrogen, 10004D) prewashed with 0.1% w/v BSA in PBS, then washed 5x with RIPA wash buffer (50 mM HEPES-KOH pH 7.5, 500 mM LiCl, 1 mM EDTA, 1% Igepal CA-630, 0.7% w/v sodium deoxycholate), once with 50 mM Tris-HCl pH 8, 10 mM EDTA, 50 mM NaCl, and eluted in elution buffer at 65°C for 30 min. Eluate was then reverse crosslinked and treated with RNase A and proteinase K, and then DNA was extracted with the ChIP DNA Clean & Concentrator-5 kit. ChIP-seq libraries were prepared using the NEBNext Ultra II DNA kit (New England Biolabs, E7645S) using up to 50 ng of input or ChIP DNA, with ∼4-8 cycles of amplification, with no pre-PCR size selection but a post-PCR double-sided 0.5x/0.9x Ampure XP bead clean-up.

### CUT&RUN

The CUTANA CUT&RUN (Epicypher) protocol and reagents (concanavalin A beads, 21-1401 and pAG MNase, 15-1016) were used with minor modifications based on the protocol from ref^101^: digestion was performed for 30 min on ice, and digestion supernatant was treated with 0.1% w/v SDS and 0.25 mg/ml proteinase K at 50°C for 1 h and then phenol-chloroform extraction was performed to extract DNA. Primary antibody incubations were performed overnight, and secondary antibody was used for mouse antibodies (TWIST1, ALX4, TCF3). Antibodies used include TWIST1 (Abcam, ab50887, 1:25), ALX4 (Novus Bio, NBP2-45490), TCF3 (Santa Cruz Biotechnology, sc-133074, 1:50), AP2a (Cell Signaling, 3215, 1:25), CTCF (Cell Signaling, 2899, 1:25), V5 (Abcam, ab206566, 1:100), Rabbit anti-mouse IgG (H+L) (Abcam, ab46540, 1:100). *E. coli* spike-in DNA (Epicypher, 18-1401) was added at 0.01 ng per reaction. Library prep was performed with modifications to the NEBNext Ultra II DNA kit as in dx.doi.org/10.17504/protocols.io.bagaibse^102^.

### SLAM-seq and RNA-seq

Cells were harvested by TRIzol (Invitrogen, 15596018) and stored at -80°C until processing. Chloroform was added to TRIzol lysate and separated into aqueous and organic phases by centrifugation per manufacturer instructions, and then the aqueous fraction was extracted using the RNA Clean & Concentrator-5 (Zymo, R1013) with on-column DNase I digestion. RNA was checked for purity by Nanodrop and was quantified by Qubit RNA broad range assay (Invitrogen, Q10210). RNA-seq libraries were prepared using the QuantSeq 3’ mRNA-Seq Library Prep FWD kit (Lexogen, 113.96) using 500 ng of input RNA and ∼15 cycles of amplification, with unique dual indices.

For acute depletions, SLAM-seq^103^ was performed as described with minor modifications, with 4 thiouridine (100 uM) labeling of nascent transcription for the last 2 h prior to harvest. Briefly, RNA was extracted using the Direct-zol RNA miniprep kit (Zymo, R2052), modified to include 0.1 mM DTT in wash buffers and 1 mM DTT to the water for elution, with protection from light. Four ug of RNA was then alkylated with 10 mM iodoacetamide (G Biosciences, 786-078) dissolved in ethanol at 100 mM, in 50% v/v DMSO, 50 mM NaPO_4_ pH 8 for 15 min at 50°C. Alkylation was quenched by addition of 1 ul of 1 M DTT, and alkylated RNA was extracted by RNA Clean & Concentrator-5 kit.

### Sequencing

Illumina sequencing libraries were sequenced using 150 bp paired-end reads on the NovaSeq 6000 and HiSeq X Ten platforms.

### Animal procedures

CD-1 mice were obtained from Charles River Laboratories and housed in RAFII facility at Stanford University. Animal care and all procedures were conducted in accordance with the Stanford University Administrative Panel on Laboratory Animal Care (under pre-approved protocol [APLAC-30364](https://login.stanford.edu/idp/profile/SAML2/Redirect/SSO?execution=e1s2)). For timed pregnancies, a female CD-1 mouse was introduced to a cage with a single CD-1 male and monitored for plugs. The noon of the day that a vaginal plug was detected was considered E0.5. Pregnant mice were sacrificed at E10.5 for dissections of facial prominences and limb buds.

### Embryo dissection

Frontonasal prominences (FNP), maxillary prominences (Mx), mandibular prominences (Md), forelimb buds (FL), and hindlimb buds (HL) of E10.5 mouse embryos were microdissected in cold PBS, and then washed twice with cold PBS, and treated with 0.05% trypsin-EDTA (Gibco, 25300054) at 37°C for 30 min, shaking at 750 rpm. Trypsin was quenched by addition of FBS, then cells were dissociated to single cells by pipetting with a P1000 pipette, chilled on ice, washed twice in PBS, and filtered through a 35-um strainer. An aliquot was taken to count cells using a Countess II, and the remainder was crosslinked and processed for ChIP as described above. One litter of embryos was used per experiment, yielding ∼1.8-3.6 million cells per region.

### Immunoprecipitation-mass spectrometry

hCNCCs were grown in 6×10-cm plates per condition and replicate, optionally treated with 500 nM dTAG^V^-1 for 30 min. Media was replaced with ice-cold PBS with 0.5 mM PMSF, cells were collected by scraping, and centrifuged at 300g for 5 min at 4°C. After aspirating supernatant, cell pellet was flash frozen and stored at -80°C. The day prior to performing IPs, 30 ug of V5 antibody (Abcam, ab206566, RRID:AB_2819156) and 6 mg magnetic beads (per sample) were conjugated overnight using the Dynabeads Antibody Coupling Kit (Invitrogen, 14311D). The next day, Dignam nuclear extraction was performed (all steps at 4°C or on ice). Briefly, cells were thawed in 5x volume buffer A (10 mM HEPES, 1.5 mM MgCl_2_, 10 mM KCl, 1x PIC and phosSTOP (Roche, 4906845001) freshly added), rotated for 5 min, centrifuged at 600g for 5 min, and resuspended in 2x buffer A. Cells were lysed by 15 strokes in a Dounce homogenizer with a tight pestle, and then centrifuged at 1000g for 5 min. The pellet was washed in 5x volume buffer A, then resuspended in 2x volume buffer C (20 mM HEPES, 25% v/v glycerol, 420 mM KCl, 1.5 mM MgCl_2_, 1x PIC and phosSTOP freshly added) and rotated for 30 min. After centrifuging at max speed for 15 min, the supernatant was slowly diluted in an equal volume of buffer D (20 mM HEPES, 25% v/v glycerol, 0.2% v/v Igepal CA-630, 1x PIC and phosSTOP freshly added) and then again diluted two-fold with buffer E (20 mM HEPES, 25% v/v glycerol, 150 mM KCl, 0.1% v/v Igepal CA-630, 1x PIC and phosSTOP freshly added). Precipitate was cleared by centrifugation at max speed for 10 min, and the supernatant (nuclear extract) was quantified by BCA assay and used for IPs. Nuclear extract was added to antibody-coupled beads pre-washed in PBS with 0.1% w/v BSA, rotated for 2h, washed four times with buffer F (20 mM HEPES, 25% v/v glycerol, 150 mM KCl, 1x PIC and phosSTOP freshly added) and two times with PBS.

In a typical mass spectrometry experiment, beads were resuspended in TEAB prior to reduction in 10 mM DTT. Reduced proteins on beads then alkylated using 30 mM acrylamide to cap cysteine residues. Digestion was performed using Trypsin/LysC (Promega) in the presence of 0.02% ProteaseMax (Promega) overnight. Following digestion and quench, eluted peptides were desalted, dried, and reconstituted in 2% aqueous acetonitrile prior to analysis.

Mass spectrometry (MS) experiments were performed using liquid chromatography (LC) using an Acquity M-Class UPLC (Waters), connect to either an Orbitrap Q Exactive HF-X (RRID:SCR_018703 Thermo Scientific) or an Orbitrap Exploris 480 (RRID:SCR_022215 Thermo Scientific). For LC separations, a flow rate of 300 nL/min was used, where mobile phase A was 0.2% (v/v) formic acid in water and mobile phase B was 0.2% (v/v) formic acid in acetonitrile. Analytical columns were prepared in-house by pulling and packing fused silica with an internal diameter of 100 microns. Columns were packed with NanoLCMS Solutions 1.9 um C18 stationary phase to a length of approximately 25 cm. Peptides were directly injected into the analytical column using a gradient (2% to 45% B, followed by a high-B wash) of 90 min. Both MS instruments were operated in a data-dependent fashion using Higher Energy Collison Dissociation (HCD).

### Protein purification, crystallization, and data collection

Expression and purification of the DNA-binding domain fragments of human TWIST1 (residues Gln101- Ser170), TCF4 (residues Arg565-Arg624), ALX4 (residues Asn210-Gln277), and BRG1 (residues 1428- 1568) were performed as described in refs ^104–106^. The DNA fragments used in crystallization were obtained as single strand oligos (Eurofins), and annealed in 20 mM HEPES (pH 7.5) containing 300 mM NaCl and 0.5 mM Tris (2-carboxyethyl) phosphine (TCEP) and 10% glycerol. The purified and concentrated proteins were mixed with a solution of annealed DNA duplex at a molar ratio 1:1:1:1.2 at room temperature, and after one hour subjected to the crystallization trials. The crystallization conditions were optimized using several conditions from JCSG crystallization kit (Molecular Dimensions). Complex was crystallized in sitting drops by vapor diffusion technique from solution containing 50 mM sodium cacodylate buffer (pH 7.5), 100 mM magnesium acetate, 18% glycerol, 20% 2-Methyl-2,4-pentanediol and 6-7% PEG (MW 8000). The X-ray data set was collected at European Synchrotron Radiation Facility (ESRF) (Grenoble, France) from a single crystal on beam-line ID23-1 at 100 K using the reservoir solution as cryo-protectant. Prior to data collection, crystals mounted on the goniometer were located and characterized using X-ray mesh scans analyzed by Dozor-MeshBest^107,108^. The experimental parameters for optimal data collection were designed using the program BEST^109^. Data were integrated with the program XDS^110^ and scaled with program AIMLESS as implemented in CCP4^111^. Statistics of data collection are presented in **Table S5**.

### Structure determination and refinement

The structure was solved by molecular replacement using program Phaser^112^ as implemented in Phenix^113^ and CCP4^111^ with the structure of TCF4 (PDB: 6OD3) as a search model for TCF4 and TWIST1, and NMR structure of ALX4 (PDB: 2M0C) as a search model for ALX4. After the positioning of proteins, the density of DNA was clear and the molecule was built manually using Coot^114^. However, we did not find any density for the BRG1 fragment. The rigid body refinement with REFMAC5 was followed by restrain refinement with REFMAC5^115^, as implemented in CCP4. Resulting statistics of the refinement are presented in **Table S5**. Structural alignments and figures were generated with PyMOL. The resulting structure was submitted to Protein Data Bank with ID 8OSB.

### ATAC-seq analysis

Reads were trimmed of Nextera adapter sequences and low-quality bases (-Q 10) using skewer v0.2.2^116^ and then mapped to the hg38 analysis set (human), mm39 (mouse), or galGal6 (chick) reference genome using Bowtie2 v2.4.1^117^ with the options --very-sensitive -X 2000. Reads were deduplicated with samtools v1.10^118^ markdup and uniquely mapped reads (-q 20) mapped to the main chromosomes (excluding mitochondria and unplaced contigs) were retained using samtools view. Read ends were shifted inward 5 bp (+5 bp on + strand, -5bp on – strand) for each fragment, and then MACS2^119^ was used to call peaks from shifted read ends using --shift -100 --extsize 200 -f BED --nomodel --keep-dup all --call-summits -- SPMR with -g hs for human, -g mm for mouse, and -g 1055580959 for chick data. Peaks from all hCNCC experiments were merged into a unified peak set by concatenating all significant summits, clustering peaks within 150 bp with bedtools^120^ cluster, keeping only the most significant summit (in any sample) per cluster with a p-value < 1E-20, extending by an additional 100 bp in both directions, and then merging any overlapping peaks with bedtools merge, resulting in 213,151 total peaks. The most significant summit within each merged peak was used as the overall summit, which was used to generate heatmaps and perform motif analyses.

Counts of reads in each sample overlapping the merged peak set were generated using bedtools, and differentially accessible peaks were called using DESeq2^121^ using only samples pertinent to each comparison, and using CNCC differentiation replicate as a covariate, excluding peaks with fewer than an average of 10 reads per dataset in the comparison. Genome browser tracks were generated by MACS2 v2.2.7.1^119^ and plotted using IGV v2.7.2^122^. Peaks with a summit within 500 bp of a TSS (from refGene GFF files from UCSC for hg38 and mm39, and ncbiRefSeq for galGal6) were considered promoter proximal, and the remaining peaks were considered distal.

For published data with multiple replicates, all summit files were concatenated and then summits within 100 bp were clustered with bedtools cluster, and the most significant summit in each cluster was retained.

For ENCODE data, bed narrowPeak files and metadata were downloaded on 1-18-2023 for all GRCh38 and mm10 non-control ATAC-seq (n=549) and DNase-seq (n=1781) experiments. All replicates were processed separately. Samples were annotated into tissue/cell types as follows: facial, limb, or lung if the annotation included that corresponding term; fibroblast if it included “fibroblast” or “HFF-Myc”, “BJ”, “AG09319”, ”AG09309”, “AG10803”, “GM03348”, “GM04504”, or“NIH3T3”; muscle if it included “muscle” or “gastroc”; neuroblastoma if it included “SK-N” or “BE2C”; and brain if it included “brain”, “cereb”, “front”, “nucleus”, “hippo”, “occipital”, “gyrus”, or “ceph”. Samples were annotated as pluripotent stem cells if the annotation included “iPS”, “WTC11”, “ES-“, “R1”, “H1”, “H7”, “H9”, “ZHBTc4”, “WW6”, “L1-S8R”, “NT2/D1”, or “GM23338” but not “NCI-H929” or “CH12.LX”.

### ChIP-seq and CUT&RUN analysis

Reads were trimmed, mapped, and deduplicated as described above for ATAC-seq analysis (but trimming Truseq adapter sequences), and then peaks were called with MACS2 (but with -f BAMPE --nomodel -- keep-dup all --call-summits –SPMR) and browser tracks were generated with deeptools v3.5.0^123^ bamCoverage -bs 10 --normalizeUsing RPGC --samFlagInclude 64 --samFlagExclude 8 --extendReads and plotted using IGV. For TWIST1 acute depletion samples, which included O9-1 mouse cranial neural crest cell spike-in chromatin, reads were mapped to a combined hg38 analysis set + mm39 reference genome. The fraction of reads mapping to the mouse genome was similar across all samples, so unnormalized tracks are shown for consistency.

For published single-end read data, reads were not deduplicated, and peaks were called with MACS2 with -f BAM and without --nomodel).

For defining TWIST1/AP2a-bound distal regions used as reference points for heatmap generation, merged ATAC peaks were defined as bound by TWIST1 or AP2a if the ATAC summit was within 200 bp of the ChIP summit, where the ChIP summits from multiple replicates were merged using bedtools cluster if they were within 150 bp. ATAC peaks were considered distal if they were at least 1000 bp from a TSS.

For comparisons of quantitative TWIST1 binding strength in hCNCC and HEK293 with and without ALX4, TWIST1 ChIP peaks (p < 10^-10^ for hCNCC, p < 10^-5^ for HEK293) from both conditions (+/- ALX4) from the same cell type were merged by removing peaks with a stronger peak within 100 bp, with bedtools cluster.

Putative enhancers (promoter-distal ATAC peaks with robust H3K27ac signal) were defined as ATAC peaks with a maximum of at least 10 RPGC in at least one TWIST1^FV^ or WT H3K27ac ChIP. For assessing log_2_ fold changes in H3K27ac signal, reads were counted over merged ATAC peaks using deeptools multiBamSummary -e --outRawCounts and used as counts for DESeq2.

CUT&RUN reads were mapped to a combined human (hg38 analysis set) and *E. coli* (K-12 substr. MG1655) reference genome using Bowtie2. CUT&RUN tracks of depleted (i.e. dTAG^V^-1 treated) samples were normalized to the control samples using the *E. coli* spike-in control, by multiplying by a scaling factor of (E_control_/H_control_)/(E_depleted_/H_depleted_), where E_x_ = fraction of reads mapped to E. coli in sample x, and Hx = fraction of reads mapped to human.

### SLAM-seq and RNA-seq analysis

Newly generated sequencing data (read 1 only) were trimmed of adapters and low-quality bases then poly A strings using skewer^116^ and processed using slamdunk v0.4.3^103^ with map options -n 100 -5 0 -q - ss, using the hg38 analysis set reference genome. Differentially expressed genes were called using DESeq2^121^ using only samples pertinent to each comparison, and using CNCC differentiation replicate as a covariate, excluding genes with fewer than 30 reads across datasets in the comparison.

Publicly available RNA-seq data were trimmed of adapters and low-quality bases using skewer^116^ and mapped using salmon^124^ quant --seqBias -l A to hg38_cdna and mm10_cdna pre-built indices (http://refgenomes.databio.org/). Salmon abundance files were summarized to the gene level and imported into R with the tximport^125^ package v1.20.0 with countsFromAbundance = ‘lengthScaledTPM’. When multiple replicates were available, the mean TPM of all replicates was used.

Human-mouse orthologs were downloaded from https://www.informatics.jax.org/downloads/reports/index.html#homology and only one-to-one orthologs were kept for analyses of RNA levels across cell types. The list of human TFs and their family definitions were downloaded from http://humantfs.ccbr.utoronto.ca/download.php (Full Database).

CCLE processed TPM values were downloaded, and these values for TWIST1 were plotted against the average for all homeodomain TFs with known motifs aligned to the HD portion of Coordinator.

### Motif analysis

Motifs from JASPAR 2018^126^, HOCOMOCO v11 human and mouse core binding models^127^, and HT-SELEX^11^, plus the Coordinator motif^20^ were used for scans of known motifs with meme suite v5.1.1 AME^99^. Motifs clusters from https://www.vierstra.org/resources/motif_clustering128 were used, with one manually added cluster for the Coordinator motif, into which the TWST1 motifs from HOCOMOCO were moved.

Motif alignments to Coordinator were performed with meme suite v5.1.1 TOMTOM^129^. Motifs from the same TF (counting orthologous human and mouse TFs as the same) and in the same cluster were collapsed, keeping the one with the best alignment.

Motif matches in the genome were calculated using meme suite v5.1.1 FIMO^130^ (for analyses of Coordinator, double E-box, and single E-box motifs in hg38 and mm39) using options –max-stored scores 5000000 and PWMScan^131^ (for annotating other motifs on hg38) using a p-value threshold of 0.001 and a background frequency of 0.25 for all bases. A p-value threshold of 10^-4^ was used to define motif presence for Coordinator, double E-box, single E-box, NEUROD1 (NDF1_HUMAN.H11MO.0.A) motifs, while a threshold of 10^-3^ was used for the HD dimer (ALX1_HUMAN.H11MO.0.B) and HD monomer motif (PRRX2_HUMAN.H11MO.0.C). For the HD monomer motif, instances overlapping Coordinator or HD dimer motifs were excluded.

ATAC-seq and ChIP-seq peaks were ranked by summit p-values as reported by MACS2 and summits ± 100 bp were used for AME and analyses of TF depletion-responsive ATAC peaks.

The double E-box motif was generated by using STREME^132^ *de novo* motif discovery to compare TWIST1 ChIP peaks (summits ± 100 bp) with significantly stronger vs. weaker binding in *ALX1^FV^ ALX4^-^*cells compared to WT hCNCCs.

For comparisons of quantitative TWIST1 binding strength in hCNCC and HEK293 with and without ALX4, merged TWIST1 summits were classified as Coordinator-containing if they had a Coordinator motif with p < 10^-4^ within 100 bp of the summit and the strongest Coordinator motif had a more significant p-value than the strongest double E-box motif; as double E-box-containing if they had a double E-box motif with p < 10^-4^ within 100 bp of the summit and the strongest double E-box motif had a more significant p-value than the strongest Coordinator motif; or otherwise as neither.

Motif logo plots were generated with meme suite v4.12.0 ceqlogo.

### IP-MS analysis

For data analysis, the .RAW data files were checked using Preview (Protein Metrics) to verify calibration and quality metrics. Data were processed using Byonic (Protein Metrics) to identify peptides and infer proteins. Proteins were held to a false discovery rate of 1%, using standard approaches described previously^133^ (Elias and Gygi, 2007). Known contaminants and any proteins with peptides detected in a control IP with the same V5 antibody on untagged hCNCC protein extracts were excluded.

### S-LDSC analysis

GWAS summary statistics for facial shape (full face, segment 1) were obtained from Figshare (https://doi.org/10.6084/m9.figshare.c.5089841.v1)134. Height GWAS summary statistics were downloaded from the Price laboratory website (https://alkesgroup.broadinstitute.org/UKBB/). LD scores were created for each annotation (corresponding to a set of differential or control distal ATAC-seq peaks) using the 1000G Phase 3 population reference. Each annotation’s heritability enrichment for a given trait was computed by adding the annotation to the baselineLD model and regressing against trait chi-squared statistics using HapMap3 SNPs with the stratified LD score regression package v.1.0.1^135^. Accessibility matched distal peaks were selected from peaks with a log_2_ fold change between -0.5 and 0.5 and adjusted p-value > 0.1 using the Matching package for R v4.10-8 with distance.tolerance = 0.01 and ties = F^136^.

## Supplementary Information

**Table S1. Motif clusters enriched in Dlx1/2/5 ChIP-seq peaks from embryonic mouse forebrain.**

**Table S2.** **Accession numbers of publicly available datasets.**

| Accession | Ref. | Samples |
| --- | --- | --- |
| GSE145327 | 28 | H9 hESC, early hCNCC, late hCNCC, D9Ch ATAC, RNA |
| GSE121840 | 37 | human HFF ATAC |
| GSE81736 | 38 | human BJ ATAC |
| GSE63577 | 34 | human BJ, HFF RNA |
| GSE80154 | 39 | human BE(2)-C, KELLY, and NGP ATAC, BE(2)-C and SHEP21 TWIST1 ChIP |
| GSE89413 | 30 | human BE(2)-C, KELLY, and NGP RNA |
| GSE113431 | 33 | mouse ESC, MEF ATAC, RNA |
| GSE89437 | 29 | mouse E8.5 NCCpro, E10.5 FNP, E10.5 Mx, E10.5 Md, E10.5 PA2 ATAC, RNA |
| GSE143914 | 35 | mouse E12.5 PM ATAC, RNA |
| GSE126293 | 36 | mouse E10.5 FL, E11.5 FL ATAC |
| GSE84795 | 32 | mouse E10.5 FL, E11.5 FL RNA |
| GSE123388 | 31 | mouse E10.5 FLHL ATAC, RNA |
| GSE185774 | 67 | chick HH22, mouse E11.5 FL ATAC |
| GSE124936 | 92 | mouse Dlx1/2/5 ChIP |

**Table S3. Immunoprecipitation-mass spectrometry data.**

**Table S4. Primers used in this study.**

**Table S5.** **X-ray crystallography data collection and refinement statistics.**

| **Statistic** | **Value** |
| --- | --- |
| Wavelength | 0.8856 |
| Resolution range | 43.73 - 2.9 (3.004 - 2.9) |
| Space group | P 31 2 1 |
| Unit cell | 60.589 60.589 237.289 90 90 120 |
| Total reflections | 125467 (13035) |
| Unique reflections | 11916 (1154) |
| Multiplicity | 10.5 (11.3) |
| Completeness (%) | 98.95 (99.91) |
| Mean I/sigma(I) | 10.16 (0.68) |
| Wilson B-factor | 86.92 |
| R-merge | 0.1895 (3.843) |
| R-meas | 0.1996 (4.025) |
| R-pim | 0.06141 (1.187) |
| CC1/2 | 0.999 (0.324) |
| CC* | 1 (0.699) |
| Reflections used in refinement | 11912 (1154) |
| Reflections used for R-free | 587 (51) |
| R-work | 0.2074 (0.3827) |
| R-free | 0.2772 (0.4247) |
| CC(work) | 0.982 (0.658) |
| CC(free) | 0.961 (0.082) |
| Number of non-hydrogen atoms | 2730 |
| Macromolecules | 2695 |
| Ligands | 5 |
| Solvent | 30 |
| Protein residues | 195 |
| RMS(bonds) | 0.016 |
| RMS(angles) | 2.36 |
| Ramachandran favored (%) | 83.60 |
| Ramachandran allowed (%) | 11.64 |
| Ramachandran outliers (%) | 4.76 |
| Rotamer outliers (%) | 29.05 |
| Clashscore | 20.99 |
| Average B-factor | 92.66 |
| Macromolecules | 98.58 |
| Ligands | 145.64 |
| Solvent | 83.28 |

Statistics for the highest-resolution shell are shown in parentheses.

## Supplemental Figures

**Figure S1.**
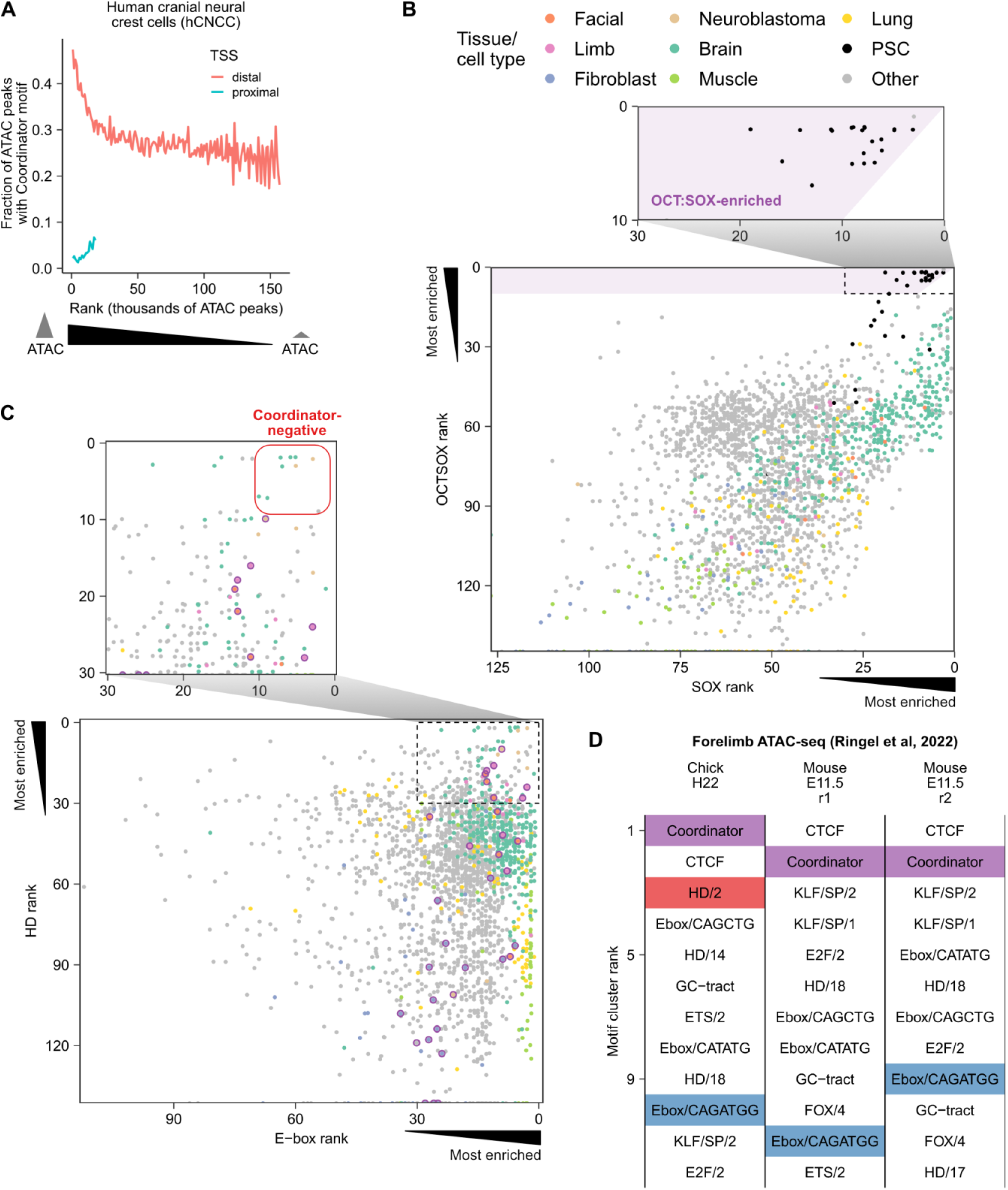
Coordinator and OCT:SOX motif enrichment in open chromatin regions, related to **Figure 1**. **A**. Coordinator motif frequency in ranked ATAC peaks ordered left to right from strongest to weakest, split by whether they overlap a TSS and grouped into bins of 1000 peaks. **B**. Rankings of OCT:SOX and its constituent SOX/1 motif in enrichment in the top 10,000 distal accessible regions, for all DNase-seq and ATAC-seq datasets on ENCODE. Points are jittered to avoid overplotting. Zoom-in highlights the pluripotent stem cell samples among those with OCT:SOX motif enrichment. **C**. Rankings of the Ebox/CAGATGG and HD/2 motif clusters in enrichment in the top 10,000 distal accessible regions, for all DNase-seq and ATAC-seq datasets on ENCODE. Points are jittered to avoid overplotting. Purple circles indicates samples with Coordinator motif enrichment (Coordinator rank < 10 and Coordinator rank < E-box and HD ranks). Zoom-in highlights samples lacking Coordinator enrichment despite enrichment of both E-box and HD motifs. **D**. Top motif clusters enriched in distal ATAC-seq peaks of chick and mouse forelimbs^67^. Coordinator is highlighted in purple, the best E-box motif match to Coordinator (Ebox/CAGATGG) in blue, and the best homeodomain motif match to Coordinator (HD/2) in red.

**Figure S2.**
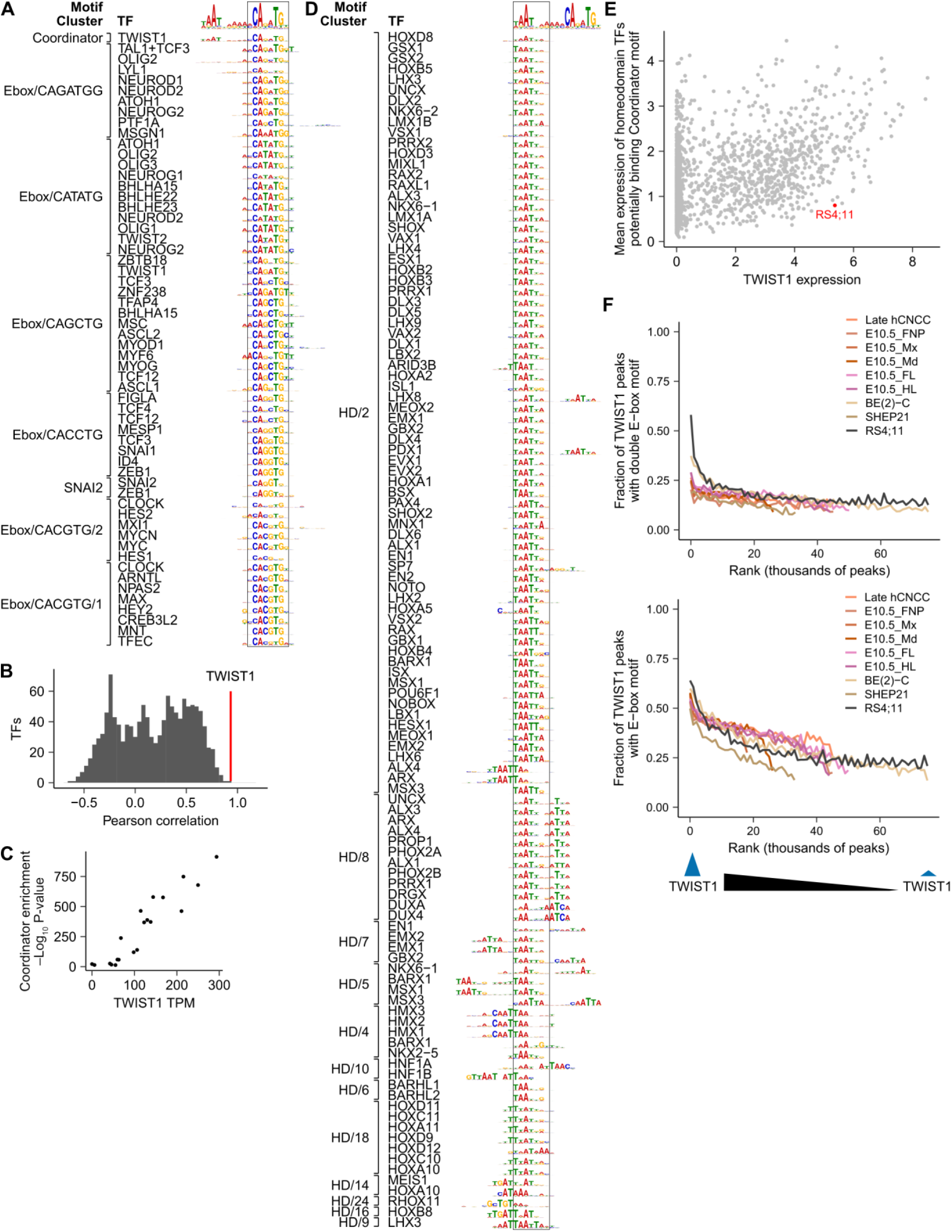
Candidate Coordinator-binding factors and a cell line without Coordinator activity, related to **Figure 2**. **A**. All bHLH and SNAI TFs with known motifs aligned to the E-box portion of Coordinator (highlighted by bounding box). **B**. TWIST1 is the TF with the highest correlation between TF RNA levels and Coordinator enrichment. **C**. TWIST1 RNA level is correlated with Coordinator motif enrichment p-value (same samples as in Figure 1D,E). **D**. All HD TFs with known motifs aligned to the HD portion of Coordinator (highlighted by bounding box). **E**. Scatter plot of TWIST1 vs average of all candidate Coordinator-binding HD TF expression in all CCLE cell lines. Both axes show log2(TPM+1) values. RS4;11 cells are highlighted in red. **F**. Frequencies of double E-box and single E-box motifs in ranked TWIST1 ChIP-seq peaks, in bins of 1000 peaks (as in Figure 2B).

**Figure S3.**
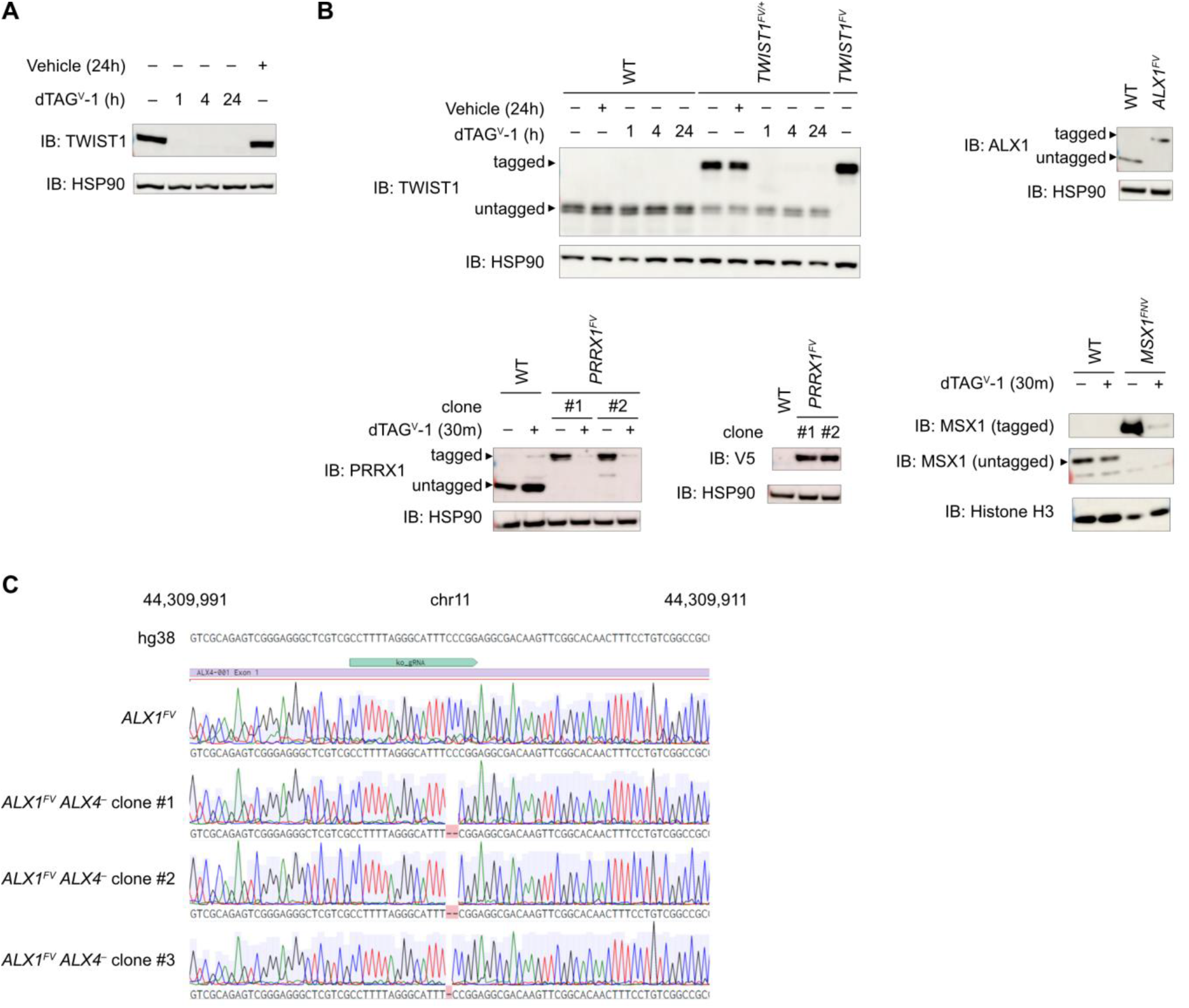
Validation of degron-tagging and ALX4 knockout, related to **Figure 3**. **A**. Western blot of TWIST1 depletion time course in *TWIST1^FV^* hCNCCs, with HSP90 as a loading control. IB: immunoblot. **B**. Western blot comparisons of tagged and untagged TF protein levels using endogenous antibodies, with HSP90 or Histone H3 as loading controls. **C**. Sanger sequencing genotyping of *ALX1^FV^ ALX4^-^* lines. The guide RNA used to generate the edits shown above traces in teal.

**Figure S4.**
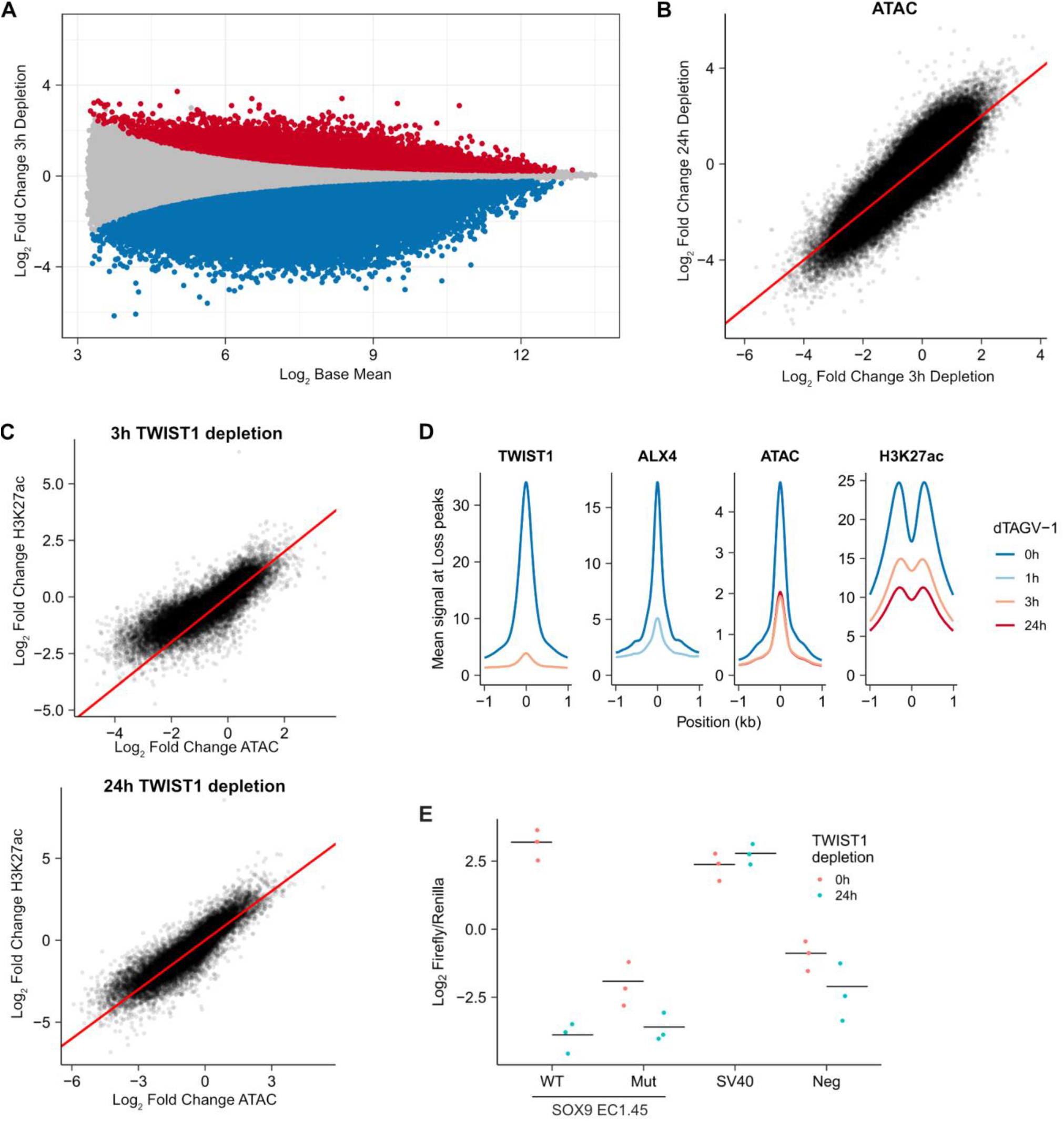
Effects of TWIST1 depletion on accessibility, H3K27ac, and enhancer activity, related to **Figure 4**. **A**. MA plot of TWIST1 3 h depletion. Significant (adjusted p-value < 0.05) upregulated and downregulated peaks are colored in red and blue, respectively. **B**. Scatter plot of 3 h vs 24 h ATAC fold changes. Red line indicates y = x. **C**. Scatter plots of ATAC vs H3K27ac fold changes upon 3 h and 24 h of TWIST1 depletion. Red line indicates y = x. **D**. Mean signal plots of TF binding, ATAC, and H3K27ac across TWIST1 depletion (dTAG^V^-1) time points (0 h to 24 h), at enhancers with loss of accessibility upon TWIST1 depletion. **E**. Luciferase enhancer reporter activity with and without TWIST1 depletion. SOX9 EC1.45 indicates the “min1-min2” enhancer from ref ^28^, Mut indicates a mutant version of the enhancer with substitutions at all high information content positions within the E-box portions of all four Coordinator motifs, SV40 is the SV40 enhancer, and Neg indicates a control plasmid lacking an enhancer insert. Points are biological replicates transfected independently (n=3).

**Figure S5.**
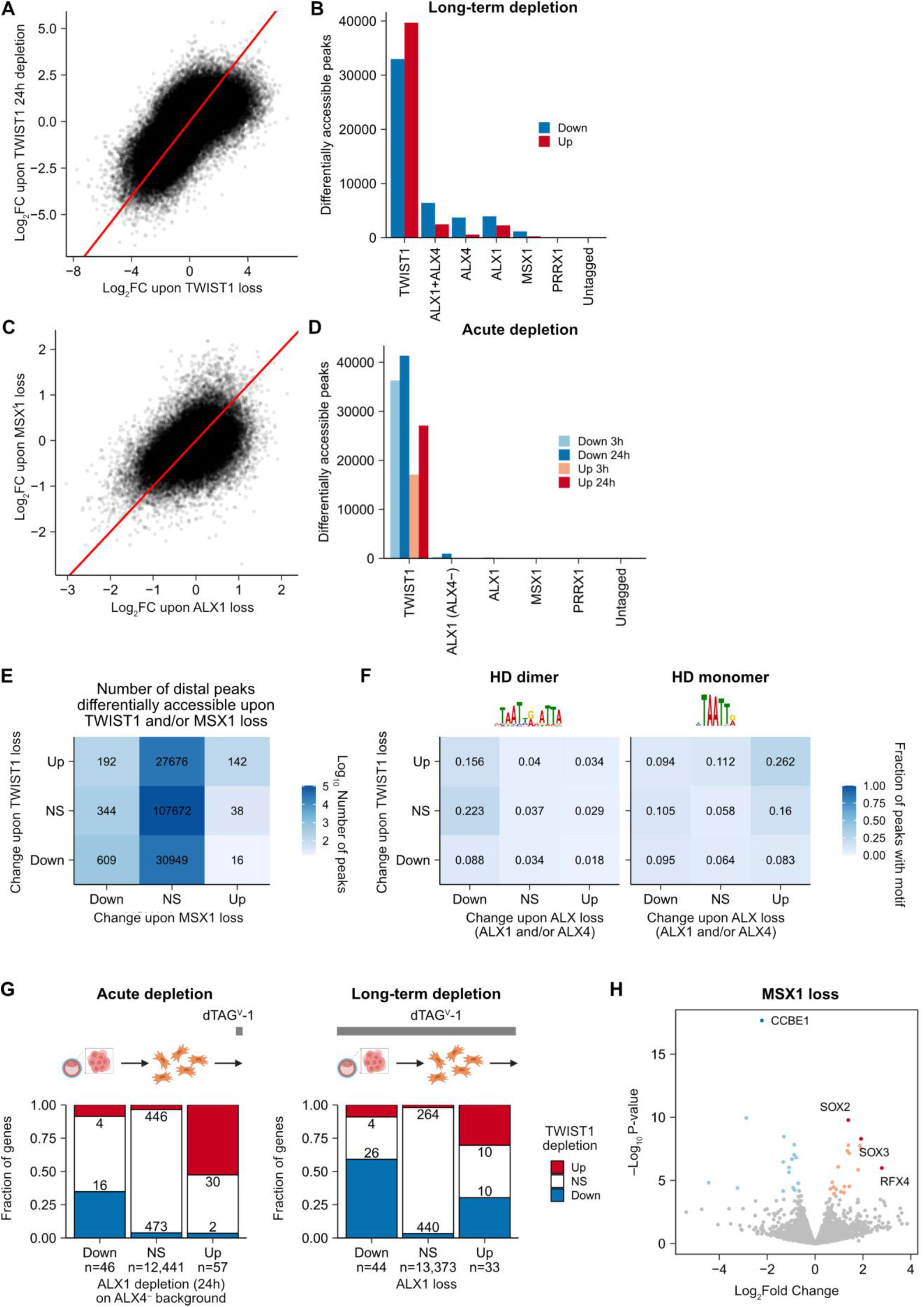
Acute and long-term depletions of Coordinator-binding TFs, related to **Figure 5**. **A**. Scatter plot of TWIST1 acute 24 h vs long-term depletion effects on accessibility at distal open chromatin. Red line indicates y = x. **B**. Bar plot of number of significant changes (FDR < 0.05) in ATAC seq upon long-term depletions. **C**. Scatter plot of ALX1 vs MSX1 long-term depletion effects on accessibility at distal open chromatin. Red line indicates y = x. **D**. Bar plot of number of significant changes (FDR < 0.05) in ATAC-seq upon acute depletions. **E**. Table of number of distal regions changing in accessibility upon MSX1 and/or TWIST1 long-term depletion. NS, not significant. **F**. Frequencies of HD motifs in regions responsive to ALX and/or TWIST1 long-term depletions. NS, not significant. **G**. Bar plots of the fraction of genes responsive to ALX1 depletion that are also responsive to TWIST1 depletion, for acute (in ALX4- background) and long-term (in ALX4+ background). NS, not significant. **H**. Volcano plot of MSX1 RNA-seq data. Significantly (FDR < 0.05) upregulated genes are highlighted in red/orange and downregulated genes are in blue. Selected genes are labeled and highlighted in darker colors.

**Figure S6.**
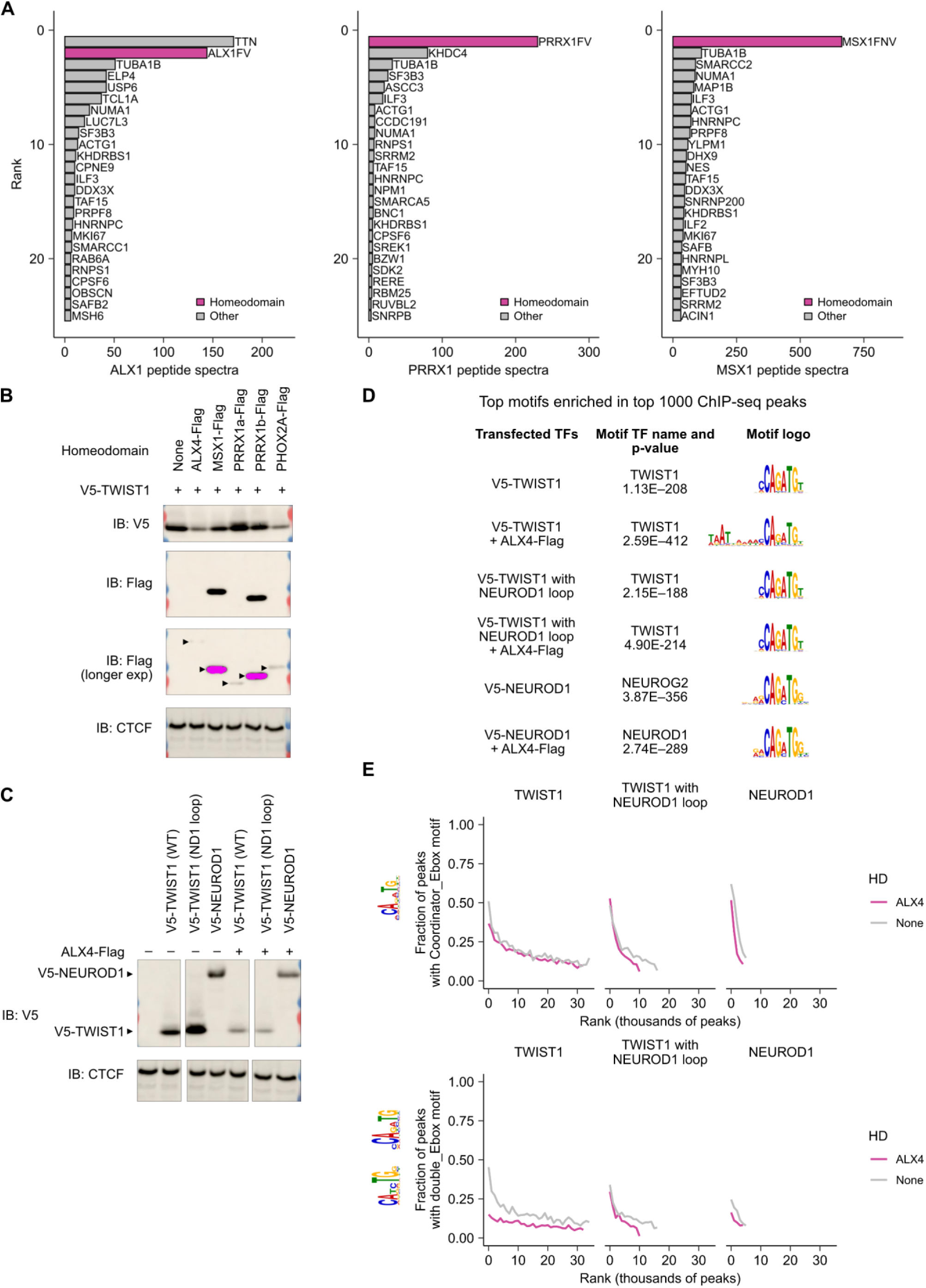
DNA guiding of TWIST1-HD interactions and variation among bHLH and HD TFs, related to **Figure 6**. **A**. Immunoprecipitation-mass spectrometry results for ALX1, PRRX1, and MSX1, using the V5 tag. The top 25 protein hits are shown, excluding known contaminants and proteins present in an untagged sample. **B**. Western blots of HEK293 cells transfected with plasmids encoding V5-TWIST1 and various homeodomain TFs, with CTCF as a loading control. IB, immunoblot. Saturated pixels are colored magenta. **C**. Western blots of HEK293 cells transfected with V5-tagged TWIST1, TWIST1 with NEUROD1 loop, or NEUROD1, each with or without ALX4-Flag. Cropped images are from same ECL reaction and exposure. **D**. Most enriched motif in the top 1000 ChIP-seq peaks for each of the six transfections shown in Figure 6G and **Figure S6C**. **E**. Motif frequency as a function of ChIP peak rank (grouped into bins of 1000 peaks), for the E-box motif within Coordinator and the double E-box motif.

**Figure S7.**
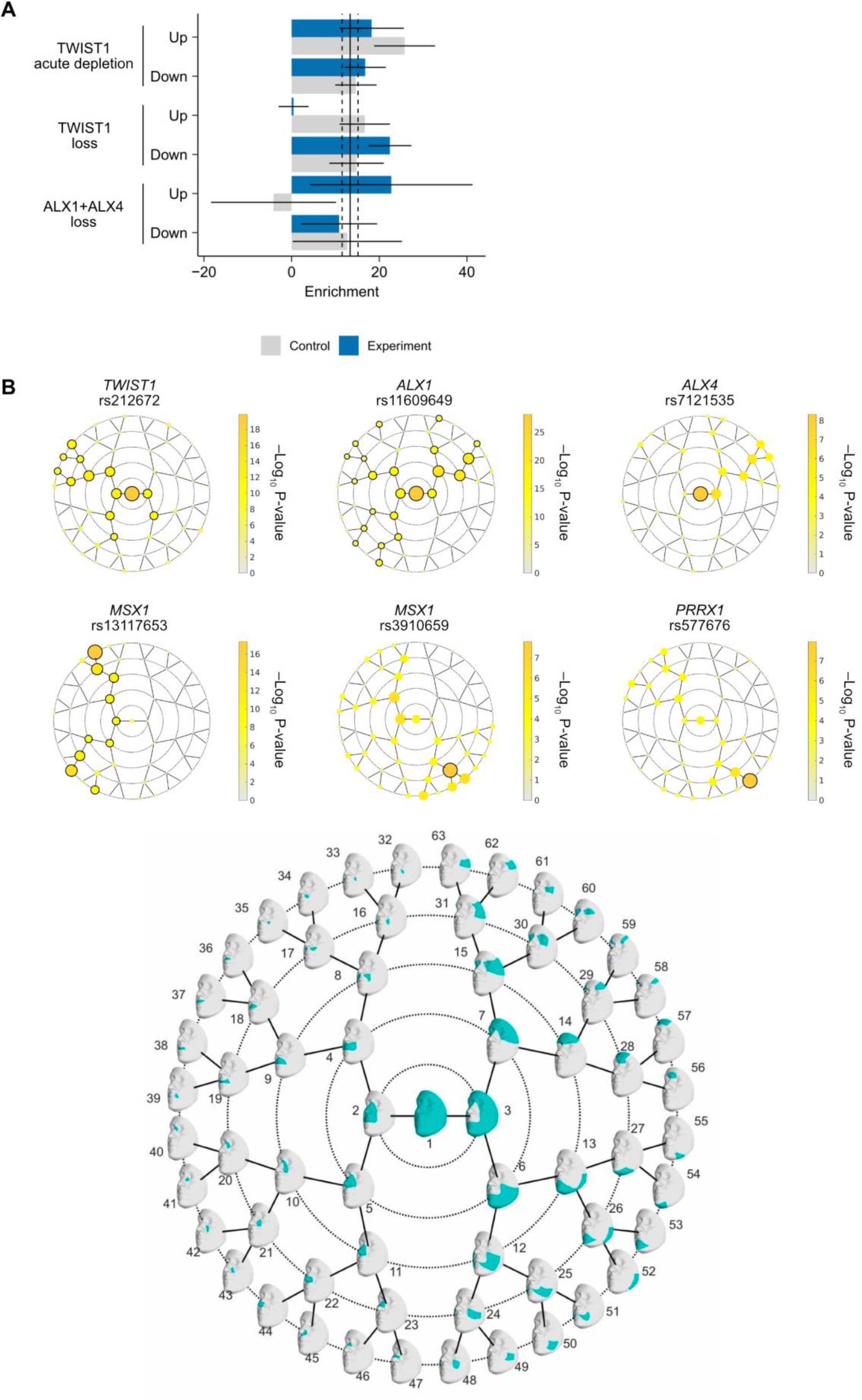
Height and facial shape heritability in Coordinator-binding TF loci, related to **Figure 7**. **A**. Fold enrichment of height-associated SNPs in distal ATAC peaks differentially accessible upon TF depletion or loss, with accessibility-matched control sets. Vertical line indicates the enrichment in all hCNCC distal ATAC peaks, with flanking dashed lines indicating error bars. Error bars represent s.e.m. **B**. Rosette plots depict the -Log^10^ P-value (one-sided, right-tailed) per facial segment in META_US_ track. Black-encircled facial segments have reached genome-wide significance (P = 5 × 10^−8^). Below, the facial segments are shown.

